# Genome-wide identification and characterization of QTLs for transcriptional noise in human midbrain cells

**DOI:** 10.1101/2025.02.01.635574

**Authors:** Naoki Hirose, Shota Mizuno, Yuki Niwa, Tomonori Hara, Hirona Yamamoto, Emiko Koyama, Junko Ueda, Takashi Tsuboi, Atsushi Takata

## Abstract

Not only the abundance of gene expression but also its cell-to-cell variation, referred to as “transcriptional noise”, is known to have certain biological significance. However, the mechanistic basis of transcriptional noise, particularly how it is regulated by genetic variants, remains elusive. In this study, we analyzed single-cell RNA sequencing (scRNA-seq) data of human induced pluripotent stem cell (iPSC)-derived midbrain cells (795,661 cells in 17 conditions) from 155 individuals with their genotypes to perform genome-wide mapping of quantitative trait loci for transcriptional noise (tnQTLs). Our analyses controlling for confounding factors such as gene expression abundance identified a total of 101,024 significant tnQTL-gene pairs. A comparison with QTLs for expression levels (i.e. eQTLs) detected by an equivalent pipeline revealed that the majority (81%) of tnQTLs were also identified as eQTLs, while no significant eQTL effects were observed in the others, and a small portion (7%) of eQTLs were with significant tnQTL effects. The tnQTLs and eQTLs showed distinctive patterns of sharing across cellular conditions, where tnQTL effects were often more condition-specific than those of eQTLs. In particular, tnQTLs without significant eQTL effects (termed tn>eQTLs) were dramatically altered by rotenone-induced oxidative stress. The tn>eQTLs also exhibited unique patterns of enrichment in various functional genomic elements, such as being frequently observed in promoters of non-QTL target genes. In the analysis using summary statistics from genome-wide association studies (GWAS) for various human complex traits, we found nominally significant enrichment of heritability for schizophrenia in tn>eQTLs. Possible contributions of tn>eQTLs to schizophrenia risk were supported by the enrichment of association signals in their target genes. We also identified genes whose transcriptional noise was implicated to be causally associated with a trait by a Mendelian randomization analysis, including *HLA* genes and *YWHAE* associated with multiple autoimmune/psychiatric disorders. To further explore the role of transcriptional noise dysregulation in disease, we analyzed scRNA-seq data from human schizophrenia and mouse model brains. Genes exhibiting differential transcriptional noise between cases and controls, i.e. differentially noisy genes (DNGs), were particularly abundant in the superficial and deep layer excitatory neurons, where association signals in schizophrenia GWAS were also enriched in their DNGs. Overall, our comprehensive mapping of tnQTLs provides a resource for a new class of regulatory genetic variants, deepens our understanding of the mechanistic basis of how genetic variants regulate transcriptional noise, and highlights the roles of tnQTLs and transcriptional noise dysregulation in human complex traits.

## Introduction

The gene expression program governs diverse biological processes, such as normal development, cellular homeostasis, response to external stimuli, and disease pathogenesis^1–4^. For the functional operation of this program, not only the steady-state amount of expressed genes but also their dynamics need to be precisely regulated. These dynamics include both continuous changes in expression and short-term fluctuations, and the magnitude of the latter can be captured as the variation in expression levels among homogeneous cell populations. This cell-to-cell variation in gene expression, referred to as “transcriptional noise”, is known to play an important role in biological events such as stochastic transitions in cellular states^5–8^, and can be quantified in a transcriptome-wide manner by analyzing single-cell RNA sequencing (scRNA-seq) data (**Figure 1A**). Indeed, the scRNA-seq technology has been utilized in studies of transcriptional noise during embryonic development, which have led to the comprehensive identification of highly variable genes, molecular characterization of them, specification of pathways involved in the regulation of transcriptional noise, and experimental demonstration of the roles of highly variable genes in differentiation^5,9–11^.

**Figure 1.**
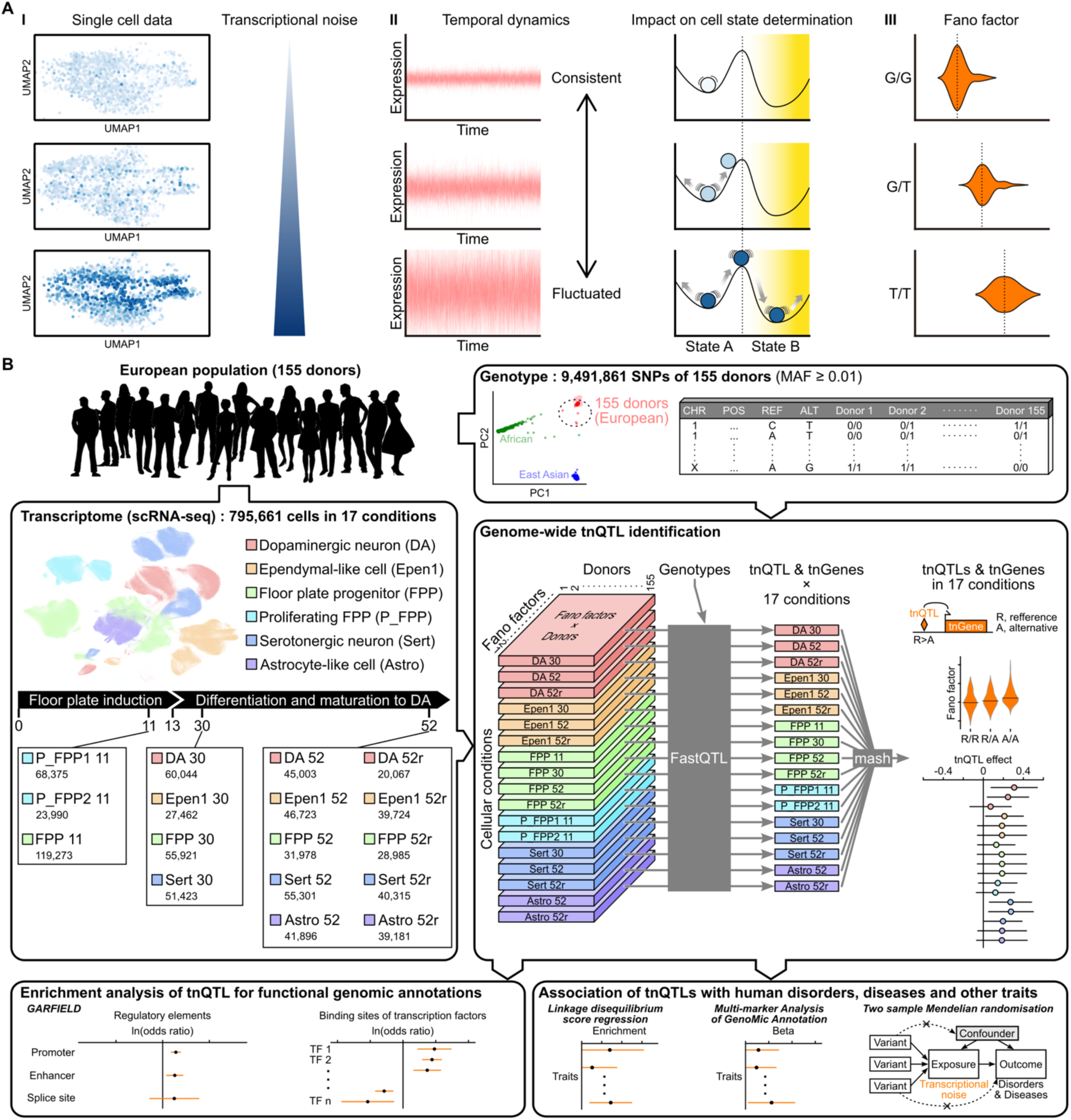
Study Overview. (A) Schematic representation of transcriptional noise quantification using scRNA-seq data. I) In cohort scRNA-seq data, the cell-to-cell variation in the expression of a gene, i.e., transcriptional noise, can vary across individuals even when the mean expression levels are the same. II) The magnitude of the transcriptional noise could reflect whether the gene exhibits consistent expression or temporal fluctuations, which is known to have certain biological significance. As illustrated by the swinging balls, the magnitude of transcriptional noise can define the chances of transitions of cells from State A to another State B, which can be a differentiated or pathological condition. III) Transcriptional noise can be represented quantitatively by the Fano factor, σ^2^/μ, where σ and μ are the standard deviation and mean of gene expression, respectively. Given the possibility that transcriptional noise can be influenced by various genomic regulatory elements characterized by specific sequences, individuals’ genotypes can be associated with its magnitude. (B) Study workflow. In this study, we used scRNA-seq data of human iPSC-derived midbrain cells from 155 donor individuals (total *n* of cells = 795,661). Cells were classified into 17 conditions based on the days *in vitro*, oxidative stress treatment (marked by “r”), and the clustering and marker gene expression patterns. Matrices of Fano factors of each gene in each donor for the 17 cellular conditions (the stacked cuboids in the figure) were separately analyzed in combination with the genotypes of 9,491,861 SNPs in the same individuals using FastQTL^25^ to identify significant tnQTL-gene pairs in each condition genome-wide. The results for each condition were then integrated using a Bayesian framework mash^26^ (see **STAR Methods** for details) to determine tnQTLs and their target genes (tnGenes) in each condition. The same analytical framework using FastQTL and mash was also applied to detect eQTLs and their target genes (eGenes).

Regarding the mechanistic basis of transcriptional noise, various components such as transcription factor (TF) binding, RNA degradation, and regulation at the protein level are supposed to play roles, and the contribution of transcriptional bursting, short windows of intense transcriptional activity interspersed with periods of inactivity, has been particularly highlighted^12–15^. Given that these components, as well as the amounts of gene expression themselves and epigenetic DNA modifications, are under the influence of genetic variants^16–21^, it is reasonable to assume that there are genomic regions associated with the levels of transcriptional noise, i.e., transcriptional noise quantitative trait loci (tnQTLs) (**Figure 1A**). Therefore, comprehensive mapping of tnQTLs at scale will enhance our understanding of the biology of transcriptional noise and their involvement in abnormal conditions, such as human diseases.

In this study, we analyze scRNA-seq data of human induced pluripotent stem cell (iPSC)-derived midbrain cells, including monoaminergic neurons, from 155 donors generated through the Human Induced Pluripotent Stem Cells Initiative (HipSci) project^22,23^ with the genotype data of the same individuals to perform genome-wide identification of tnQTLs. Our analyses controlling for confounding factors such as gene expression abundance identify a total of 101,024 significant tnQTL-gene pairs. By investigating the properties of these tnQTLs, we delineate their relationship to QTLs for gene expression levels (i.e. expression QTLs: eQTLs), dynamics across different cellular conditions, enrichment patterns in *cis*-regulatory elements, and potential roles in human complex traits, which collectively contribute to a better understanding of the biological meaning and underpinnings of transcriptional noise.

## Results

### Genome-wide identification of tnQTLs

To comprehensively identify tnQTLs, we first analyzed scRNA-seq data of iPSC-derived midbrain cells from 155 donors to quantify the transcriptional noise of each gene in each individual for the 17 cellular conditions shown in **Figure 1B** (days 30 and 52 dopaminergic neurons [DA]; days 30 and 52 ependymal-like cells [Epen1]; days 11, 30, and 52 floor plate progenitors [FPP]; day 11 proliferating FPP type 1 and 2 [P_FPP1 and P_FPP2]; days 30 and 52 serotonergic-like neurons [Sert]; day 52 astrocyte-like cells [Astro], day 52 cells include those treated with rotenone to mimic oxidative stress conditions and untreated cells) (**Supplemental Table S1**). Quantification of transcriptional noise was performed by calculating Fano factors from the normalized expression read counts (see **STAR Methods; Supplemental Figure S1**). The Fano factor is a measure of variability computed as the ratio of the variance to the mean, thus controlling for the effect of gene expression abundance on the transcriptional noise. When we evaluated the characteristics of genes with large transcriptional noise and those with stable expression by performing a Gene Set Enrichment Analysis (GSEA) using clusterProfiler^24^, we observed that the expression of organellar and mitochondrial large ribosomal subunit genes was commonly stable in undifferentiated day 11 P_FPP and differentiated day 52 DA/Sert cells, whereas cytosolic ribosomal protein genes, which are non-overlapping with organellar and mitochondrial ribosomal genes, were noisy in multiple conditions (**Supplemental Figure S2**). The obtained transcriptional noise data (the gene Fano factor × donor matrix in **Figure 1B**) for the 17 cellular conditions were then analyzed separately using FastQTL^25^ by incorporating the corresponding donor genotype data (9,491,861 single nucleotide polymorphisms [SNPs] after quality controls; **Supplemental Figure S3**) and covariates to identify tnQTLs in each condition (**Supplemental Figure S4**). Subsequently, we integrated the data of tnQTLs in each condition using Multivariate Adaptive Shrinkage (mash^26^), a statistical method that allows for the synthesis of multiple datasets by taking into account their correlation structure, to identify shared and condition-specific tnQTLs with improved statistical power (**Figure 1B**).

Through these analyses, we identified a total of 101,024 tnQTL-gene pairs (11,352 after pruning for *r*^2^ > 0.8 in a 100 kb window) with a local false sign rate (lfsr)^26,27^ < 0.05 in the integrated dataset of 17 conditions (**Figure 2A** and **Supplemental Table S2**, see **STAR Methods** for details). The lfsr is analogous to the local false discovery rate (FDR) but measures confidence in the sign of each effect in the context of multiple hypothesis testing^27^. In each condition, 53,605-78,487 tnQTL-gene pairs were detected. The total number of unique genes whose transcriptional noise was regulated by the identified tnQTLs (tnGenes) was 2,016, and the number of tnGenes in each condition ranged from 638 to 938 (**Figure 2B** and **Supplemental Table S2**). To compare with tnQTLs, we also mapped eQTLs in these cells using the same analytical procedures, identifying a total of 1,224,228 eQTL-gene pairs (**Figure 2C**) influencing the expression abundance of 7,306 genes (eGenes) (**Figure 2D**). When we assessed the overlap between tnQTLs and eQTLs, we found that 81% of tnQTL-gene pairs have an effect as an eQTL in the integrated dataset (**Figure 2E**, **Supplemental Figure S5A**, and **Supplemental Table S2**). Since we controlled for the effect of expression abundance on transcriptional noise in our exploration of tnQTLs, these were considered to have the properties of both tnQTL and eQTL. Therefore, we defined these QTLs as “tn+eQTLs”. Although the majority of tnQTLs were also identified as eQTLs, no significant eQTL effects were observed in the remaining 19% of the tnQTLs. We refer to these as “tn>eQTLs”. Reflecting the much smaller number of tnQTLs compared to eQTLs, a minor proportion of eQTL-gene pairs (7%) showed effects as tnQTLs (**Figure 2E**). We considered the other eQTLs with no significant effects as tnQTLs to be “e>tnQTLs”. In total, there were 18,794 tn>eQTLs, 82,230 tn+eQTLs, and 1,141,998 e>tnQTLs (**Figure 2E**). Similarly, we analyzed the overlap between tnGenes and eGenes and found 681 tn>eGenes, 1,335 tn+eGenes, and 5,971 e>tnGenes in the integrated dataset (**Figure 2F** and **Supplemental Figure S5B**).

**Figure 2.**
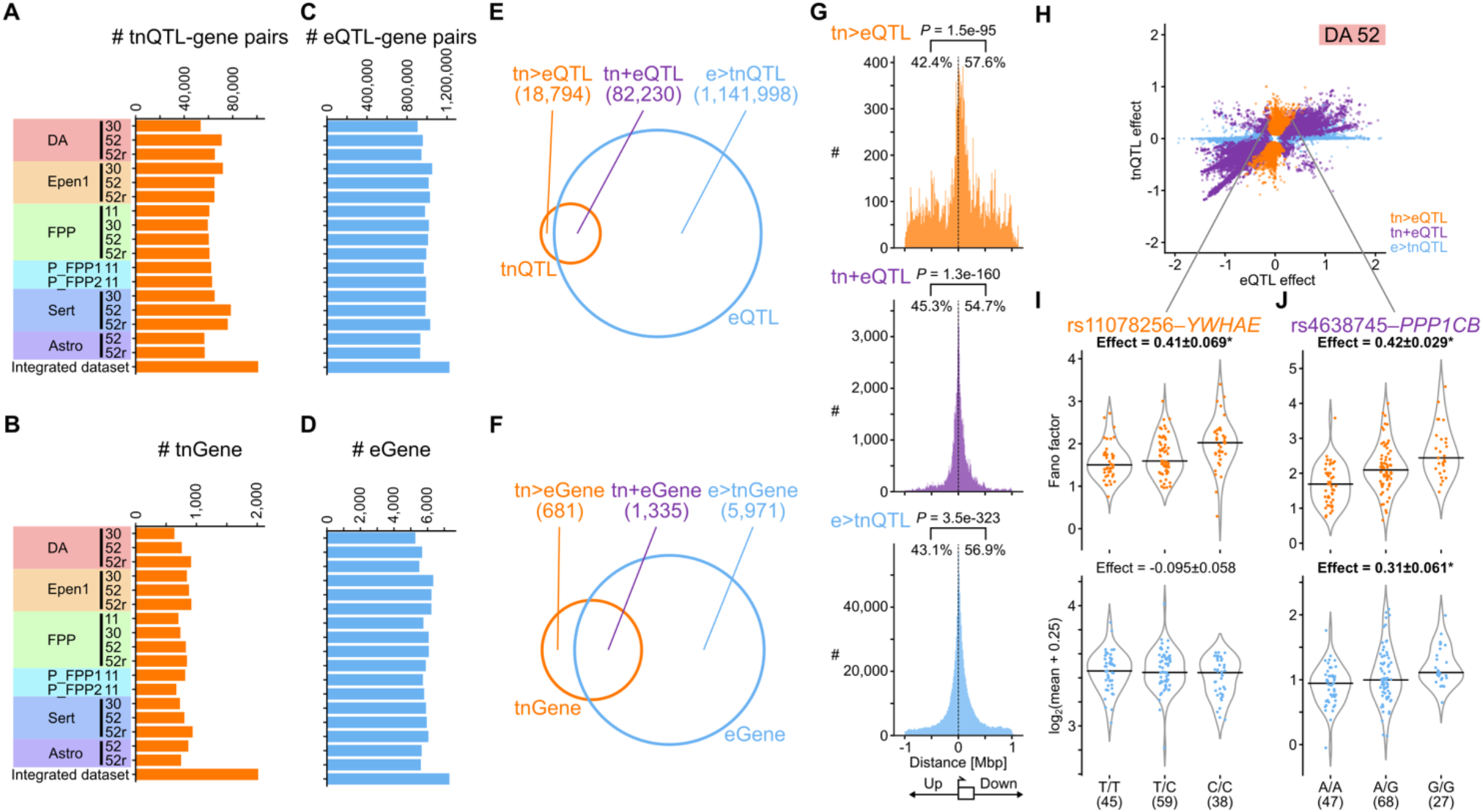
Genome-wide identification of tnQTLs. (A-D) Bar plots of the numbers of tnQTL-gene pairs (A), tnGenes (B), eQTL-gene pairs (C), and eGenes (D) in the integrated dataset and each condition. (E and F) Venn diagrams showing the overlap between tnQTLs and eQTLs (E) and tnGenes and eGenes (F) determined in the integrated dataset. The numbers of tn>eQTLs, tn+eQTLs, and e>tnQTLs (E) and tn>eGenes, tn+eGenes and e>tnGenes (F) are indicated in the parentheses. (G) Distributions of the distances to the TSS of the target genes from tn>eQTL (top), tn+eQTL (middle), and e>tnQTL (bottom) SNPs. Distances were calculated in a way aware of the transcriptional direction. The proportions of QTLs upstream and downstream of the TSS are shown. *P* values were calculated by two-tailed binomial tests. (H) A scatter plot of the effect sizes (the posterior mean of effects estimated by mash) of tnQTLs and eQTLs. The result for day 52 DA is shown. The tn>eQTLs, tn+eQTLs, and e>tnQTLs in the integrated dataset are indicated by orange, purple, and cyan circles, respectively, and are overlaid in this order from the foreground. Note that the three types of QTLs were defined based on the results in the integrated dataset, and some of them are not significant in the day 52 DA cells. (I and J) Violin plots of the Fano factors (top) and mean expression levels (bottom) of representative tn>eQTL (I, rs11078256 targeting *YWHAE*) and tn+eQTL (J, rs4638745 targeting *PPP1CB*) per genotype. The posterior mean of estimated effect ± the standard deviation for each SNP is shown above the plot, and significant effects (lfsr < 0.05) are in bold and marked with an asterisk. The horizontal black lines indicate the median.

When we assessed the distributions of the distances to the transcription start site (TSS) of their target genes from tn>eQTL, tn+eQTL, and e>tnQTL SNPs, as expected, all types of QTLs were concentrated around the TSS (**Figure 2G**), indicating that QTLs are correctly called in this study. Also, our analysis accounting for the direction of transcription showed that tn>eQTL, tn+eQTL, and e>tnQTL were all more abundant downstream of the TSS. **Figure 2H** shows a scatter plot of the effect sizes of each SNP as a tnQTL (y-axis) and as an eQTL (x-axis) in the day 52 DA cells. Since the effect sizes of tnQTLs were generally smaller than those of eQTLs, it was reasonable that we identified fewer significant tnQTLs in this study. Transcriptional noise and expression abundance for each genotype of a representative tn>eQTL (rs11078256 targeting *YWHAE*) and a tn+eQTL (rs4638745 targeting *PPP1CB*) are highlighted in **Figures 2I** and **J**, respectively.

### Similarities and divergences of tnQTLs and eQTLs across cellular conditions

Taking advantage of our comprehensive datasets of tnQTLs and eQTLs in 17 different midbrain cell types, we then analyzed the similarities and divergences of these QTLs across cellular conditions. When we evaluated how many conditions significant QTL effects were shared, we found that more than half (50.3%) of e>tnQTLs were called in all cellular conditions in our mash analysis (**Figure 3A**, right), whereas the proportion of tn>eQTLs common to all conditions was lower (32.4%) and 25.4% were identified in only one cell type (**Figure 3A**, left). Furthermore, less overlap of significant tnQTL effects across cellular conditions was supported by an analysis of tnQTL and eQTL effects in a single dataset of tn+eQTLs (**Figure 3A**, middle two), where the number of SNPs with significant tnQTL effects in all conditions was fewer (25.2% vs. 32.7%) and more SNPs exhibited significant effects only in one condition (4.2% vs. 0.8%). Considering that these analyses based on the overlap of significant QTLs across conditions would be potentially biased by the difference in detection power between tn>eQTLs and e>tnQTLs, we then performed an analysis of correlations of QTL effect sizes (**Figure 3B**). We again found that the tnQTL effects were less correlated than those of eQTLs. We also confirmed that these less overlap and correlation of the tnQTL effects are not explained by the expression specificities of QTL target genes, as indicated by the plots of the maximum and median expression levels of each gene across the 17 cellular conditions (**Supplemental Figure S6**). While it is expected that the maximum and median expression levels of genes specific to certain conditions would be non-zero and zero, respectively^28^, the proportion of such genes, which are less likely to overlap or correlate across conditions, in tn>eGene was rather smaller than the other types of genes.

**Figure 3.**
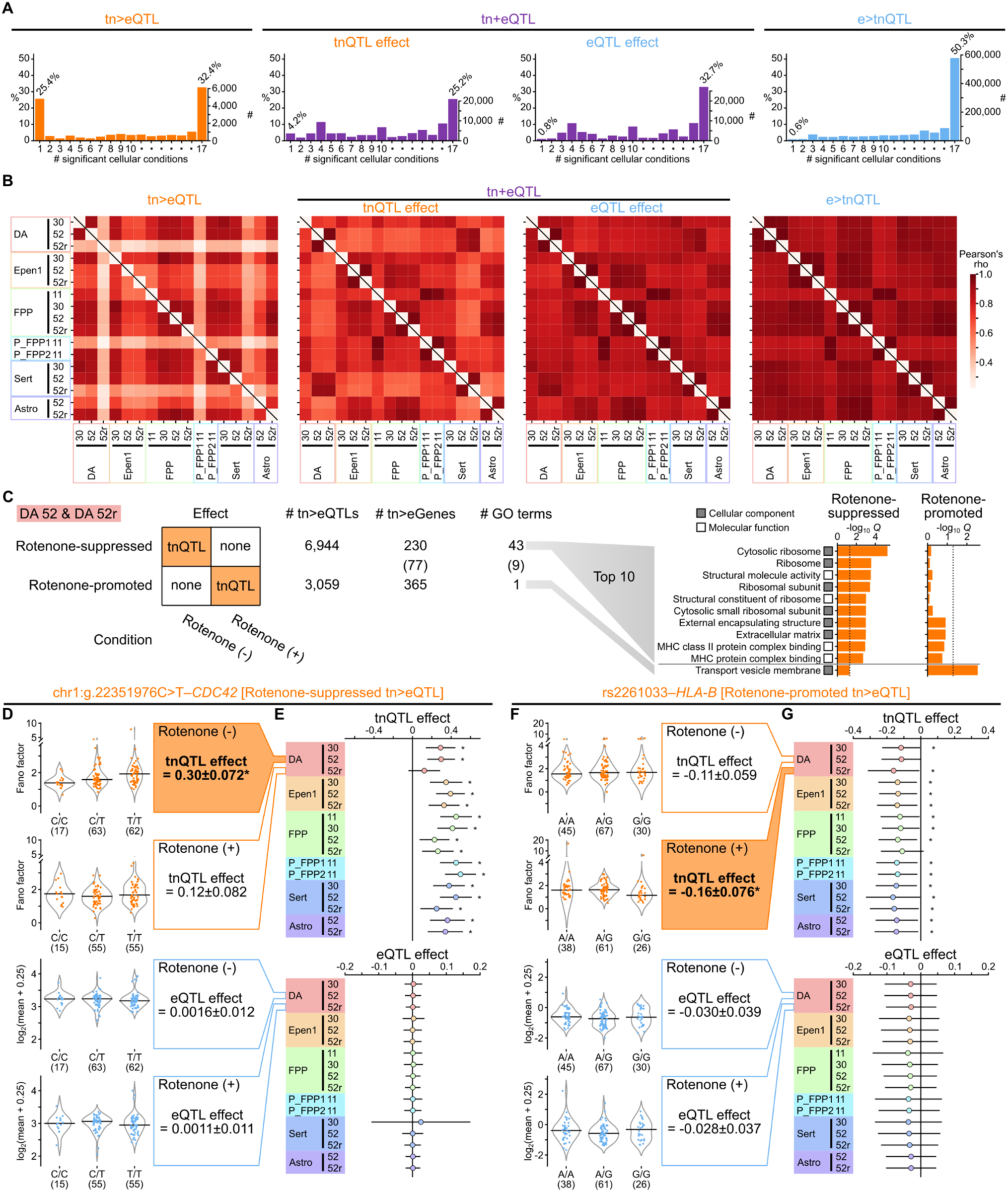
tnQTLs and eQTLs across cellular conditions. (A) Distributions of the numbers of cellular conditions where significant QTL effects were shared. From left to right, results for significant tn>eQTLs, tnQTL effects of tn+eQTLs, eQTL effects of tn+eQTLs, and e>tnQTLs in the integrated dataset are shown. The x-axis indicates the number of cellular conditions in which we observed significant QTL effects, with 1 and 17 denoting that the effect was significant only in a single condition and all individual conditions, respectively. The y-axis shows the proportions (left scale) and the numbers (right scale) of QTLs corresponding to each number of significant conditions (1-17), with the proportions for “1” and “17” displayed above the bars. (B) Correlation heatmaps of tnQTL effects of tn>eQTLs, tnQTL effects of tn+eQTLs, eQTL effects of tn+eQTLs, and eQTL effects of e>tnQTLs across 17 cellular conditions. We used the data of QTL effect sizes in each condition for QTLs that were significant in the integrated dataset. (C) Left, The numbers of tn>eQTLs that showed significant effects only in rotenone-untreated day 52 DA/Sert cells (rotenone-suppressed) and those significant only in rotenone-treated cells (rotenone-promoted), their target tn>eGenes, and GO terms enriched among these genes. The numbers of tn>eGenes targeted by both rotenone-suppressed and promoted tn>eQTLs and the GO terms commonly detected in the analyses of rotenone-suppressed and promoted tn>eQTL target genes are shown in the parentheses. Right, Top ten GO terms significantly enriched among tn>eGenes targeted by rotenone-suppressed tn>eQTLs and a term significant among tn>eGenes targeted by rotenone-promoted tn>eQTLs. The bars indicate -log_10_ *Q* values for enrichment. The gray and white squares correspond to the GO categories (cellular component and molecular function). (D-G) Violin plots of the Fano factors (top) and mean expression levels (bottom) of representative rotenone-suppressed (D) and rotenone-promoted (F) tn>eQTLs SNP-gene pairs in day 52 DA cells. The posterior mean of estimated effect ± the standard deviation for each SNP is shown, and significant effects (lfsr < 0.05) are in bold and marked with an asterisk. The horizontal black lines indicate the median. Panels E and G show the posterior mean of estimated tnQTL (top) and eQTL (bottom) effects ± 2 × the standard deviation across the 17 cellular conditions. *lfsr < 0.05.

Interestingly, our analysis of the correlations between rotenone-treated day 52 DA/Sert cells and other cell types showed that only the tnQTL effects of tn>eQTLs were poorly correlated (**Figure 3B**, left), suggesting that this type of effects are dynamically regulated in response to oxidative stress. By extracting tn>eQTLs that showed significant effects only in rotenone-untreated or treated cells, we identified 6,944 and 3,059 tn>eQTLs that were suppressed and promoted by the exposure to oxidative stress in day 52 DA cells, respectively (**Figure 3C**, left). These rotenone-suppressed and promoted tn>eQTLs respectively targeted 307 and 442 tn>eGenes (77 in common), and among these tn>eGene sets 52 and 10 GO terms (9 in common) were significantly enriched. Thus, while there were fewer rotenone-promoted tn>eQTL SNP-tnGene pairs in day 52 DA cells, they targeted a larger number of genes, yet these genes converged on specific biological categories as indicated by a smaller number of significantly enriched GO terms. This pattern was confirmed in the analysis of day 52 Sert cells (**Supplemental Figure S7B** and **D**). GO terms that were specifically enriched in rotenone-suppressed tn>eGenes in day 52 DA cells included those related to cytosolic ribosome, structural molecular activity, and major histocompatibility complex (MHC) protein ligands, whereas transport vesicle membrane genes were enriched in rotenone-promoted tn>eGenes (**Figure 3C**, right, and **Supplemental Figure S7A** and **C**). In **Figures 3D-G**, the tnQTL and eQTL effects of representative rotenone-suppressed (chr1:g.22351976C>T-*CDC42*, **Figures 3D** and **E**) and rotenone-promoted (rs2261033-*HLA-B*, **Figures 3F** and **G**) tn>eQTL SNP-gene pairs in day 52 DA cells (**Figures 3D** and **F**) and 17 cellular conditions (**Figures 3E** and **G**) are exemplified.

### Enrichment of tn>eQTLs, tn+eQTLs, and e>tnQTLs in functional genomic elements

Identification of functional genomic elements enriched for tnQTLs can provide insight into the regulatory mechanisms of transcriptional noise. To this end, we analyzed the enrichment of tn>eQTLs, tn+eQTLs, and e>tnQTLs in various features of genomic intervals using GARFIELD^29^, a statistical method controlling for key confounding factors such as distance to a TSS and linkage disequilibrium.

When we tested enrichment in a total of 86 genomic elements defined based on annotations in FANTOM5^28,30,31^, ENCODE^32^, GENCODE^33^, G-quadruplex-sequencing^34^ and RepeatMasker^35^, we found that three types of QTLs were commonly enriched in various elements such as transcribed enhancers and *Alu* elements (**Figure 4A**, **Supplemental Figure S8**, and **Supplemental Table S3**). In many of these elements, tn+eQTLs showed the largest effect sizes, indicating that SNPs hitting known regulatory elements were more likely to be associated with both quantitative traits than those in functionally uncharacterized regions. On the other hand, only the tn>eQTL did not show significant enrichment in several elements, including TATA-containing promoters and splice sites, with smaller effect sizes compared to the other QTL types (**Figure 4A**). Regarding QTL enrichment in promoter regions, we further performed an analysis classifying QTL SNPs into those that are located in promoters immediately upstream of their target genes (“IN” in **Figure 4B**, top) and the others, that is, those present upstream of non-target genes (“OUT” in **Figure 4B**, top). In all of the analyses based on the four promoter definitions (TATA & CpG, TATA & CpG-less, TATA-less & CpG, and TATA-less & CpG-less), we observed that the proportions of “IN” QTLs were lowest in tn>eQTLs (**Figure 4B**, bottom). This may suggest that the effects of tn>eQTLs in promoters are not mediated by regulation of the immediate downstream genes but rather are explained by other functions, such as distal enhancer activity of promoters^36–38^.

**Figure 4.**
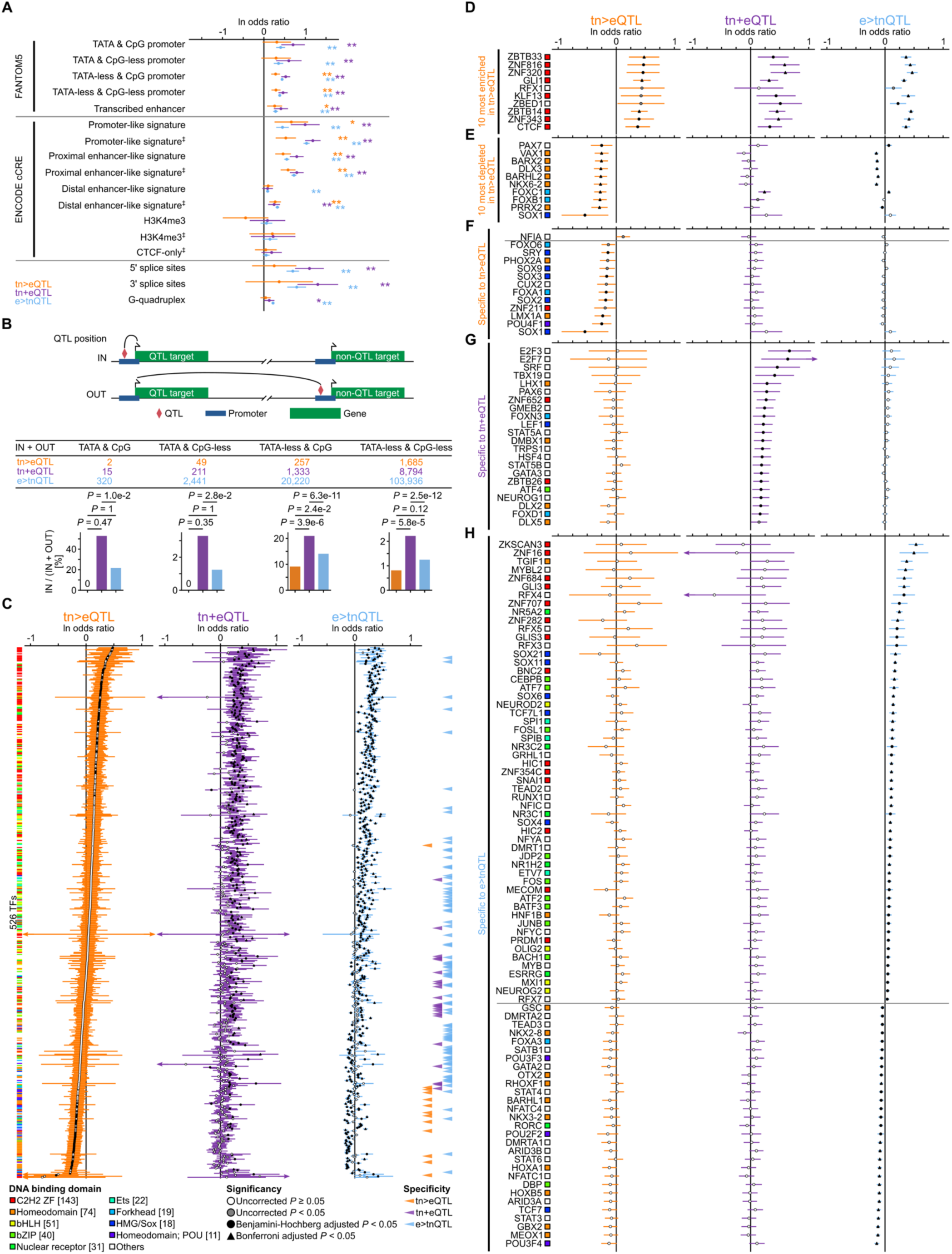
Enrichment in functional genomic elements. (A) Enrichment of tn>eQTLs (orange), tn+eQTLs (purple), and e>tnQTLs (cyan) in various regulatory elements and genomic features evaluated by GARFIELD^29^. ‡, CTCF-bound. \**P* < 0.05; \*\**P* < 0.05/51 (i.e., significant after Bonferroni correction). (B) Top, schematic representation of the relationship between the location of a QTL and its target gene. We defined QTL SNPs in promoters immediately upstream of their target genes as “IN”, and the other promoter QTL SNPs present upstream of non-target genes to be “OUT”. Middle, a table of the numbers of tn>eQTLs, tn+eQTLs, and e>tnQTLs within the four types of the FANTOM5 promoters (total numbers of IN and OUT). Bottom, Bar plots showing the proportions of tn>eQTLs (orange), tn+eQTLs (purple), and e>tnQTLs (cyan) within a promoter of their target genes (IN) among all promoter QTLs (IN+OUT). *P* values were calculated by two-tailed Fisher’s exact tests. (C) Enrichment of tn>eQTLs, tn+eQTLs, and e>tnQTLs in the predicted TF binding sites, sorted in the order of enrichment effect sizes for tn>eQTLs. TFs whose binding sites are specifically enriched or depleted for tn>eQTLs, tn+eQTLs, and e>tnQTLs are indicated by the orange, purple, and cyan arrowheads, respectively. (D and E) Top ten TFs whose binding sites are most enriched (D) or depleted (E) for tn>eQTLs. (F-H) TFs whose binding sites are specifically enriched or depleted for tn>eQTLs (F), tn+eQTLs (G), and e>tnQTLs (H). In (A) and (C-H), the error bars indicate 95% confidence intervals. In (C-H), the black triangles, black circles, gray circles, and white circles indicate those with Bonferroni-corrected *P* < 0.05, Benjamini-Hochberg adjusted *P* < 0.05, uncorrected *P* < 0.05, and uncorrected *P* ≥ 0.05, respectively. TFs are color-coded by their DNA binding domains as shown at the bottom of (C); the numbers of TFs with each domain are indicated in the square brackets. andard deviation across the 17 cellular conditions. *lfsr < 0.05.

Next, we examined enrichment of tn>eQTLs, tn+eQTLs, and e>tnQTLs in TF binding sites predicted by JASPAR (**Figure 4C**, and **Supplemental Table S3**). Of the 526 TFs whose expression was detected in the integrated dataset, 141 and 81 TFs showed enrichment and depletion of tn>eQTLs in their TF binding sites at uncorrected *P* < 0.05, respectively. Eight of the ten TFs with the largest effect sizes in tn>eQTL enrichment were those with C2H2 Zinc finger domains and many of them also exhibited significant enrichment of tn+eQTLs and e>tnQTLs (**Figure 4D**). Among the TFs with tn>eQTL depletion, the smallest effect size was observed for SOX1. Several of these TFs, including FOXC1, showed a contradictory pattern with tn>eQTLs depletion and enrichment for other types of QTLs (**Figure 4E**). We found that 13, 22, and 84 TFs with enrichment or depletion specific to tn>eQTLs, tn+eQTLs, and e>tnQTLs, respectively (uncorrected *P* < 0.05 in one type and ≥ 0.05 in the others, **Figures 4F-H**). Among the TFs specific to tn>eQTLs, NFIA showed enrichment, whereas depletion was observed for the other 12 TFs. The pattern that there are more TFs with depletions was characteristic of tn>eQTLs and not seen in tn+eQTLs or e>tnQTLs (**Figures 4G-H**).

Depletion of tn>eQTLs could be observed when fewer SNPs in the binding sites of a given TF are associated with this quantitative trait than would be expected by random chance, but may also be explained by the overall negative selection against tn>eQTLs. Although it is difficult to clarify which of these two possibilities is closer to the truth, our analysis showed that tn>eQTL SNPs have lower derived allele frequencies than tn+eQTL, e>tnQTL, and the other non-QTL SNPs (**Supplemental Figure 9A**). Also, the bases hit by tn>eQTL SNPs are more evolutionarily conserved than those hit by tn+eQTL or e>tnQTL SNPs (**Supplemental Figure 9B**). Therefore, tn>eQTLs may be more negatively selected than tn+eQTLs and e>tnQTLs, or tn+eQTLs and e>tnQTLs would be positively selected. In line with this, the TFs whose binding sites were depleted of the midbrain cell tn>eQTLs identified in this study were found to be highly interconnected (**Supplemental Figure 9C**) and significantly enriched for genes involved in essential biological processes related to brain development and differentiation, such as neuron differentiation (GO:0030182) and neurogenesis (GO:0022008), in our GO enrichment analysis using all expressed TFs as the background geneset (**Supplemental Figure 9C**). In addition, tn>eGenes regulated by tn>eQTLs significantly overlapped with human constrained genes with less loss-of-function (LoF) variants than expected (loss-of-function observed/expected upper bound fraction [LOEUF] score < 0.6), while tn+eGenes and e>tnGenes showed significantly less overlap than expectation (**Supplemental Figure 9D**).

Taken together, these results suggest that QTLs that specifically affect transcriptional noise (i.e. tn>eQTL) and QTLs for expression levels (tn+eQTL and e>tnQTL) have different enrichment and depletion patterns of functional genomic elements and target different characteristics of genes in terms of evolutionary conservation and constraint.

### Enrichment of tnQTLs in genes and loci associated with various human complex traits

Previous studies have consistently demonstrated that QTLs for various molecular phenotypes contribute to human diseases and traits. In this context, we evaluated how tn>eQTLs, tn+eQTLs, and e>tnQTLs and their target genes are enriched in loci associated with human complex traits using two statistical approaches, Linkage Disequilibrium SCore regression (LDSC^39,40^) and Multi-marker Analysis of GenoMic Annotation (MAGMA^41^). Both of these methods take into account confounding factors such as linkage disequilibrium and gene size. In the LDSC analysis, which allows SNP lists to be used as inputs, we evaluated if tn>eQTL, tn+eQTL, and e>tnQTL SNPs were enriched in trait-associated loci explaining the heritability, and in the MAGMA analysis, which uses gene lists as inputs, we analyzed if the sets of tn>eGenes, tn+eGenes, and e>tnGenes were in aggregate associated with traits. Considering that this study uses data derived from midbrain cells, we analyzed the 21 traits in **Figure 5**, which included a variety of brain-related phenotypes (see **STAR Methods** for details).

**Figure 5.**
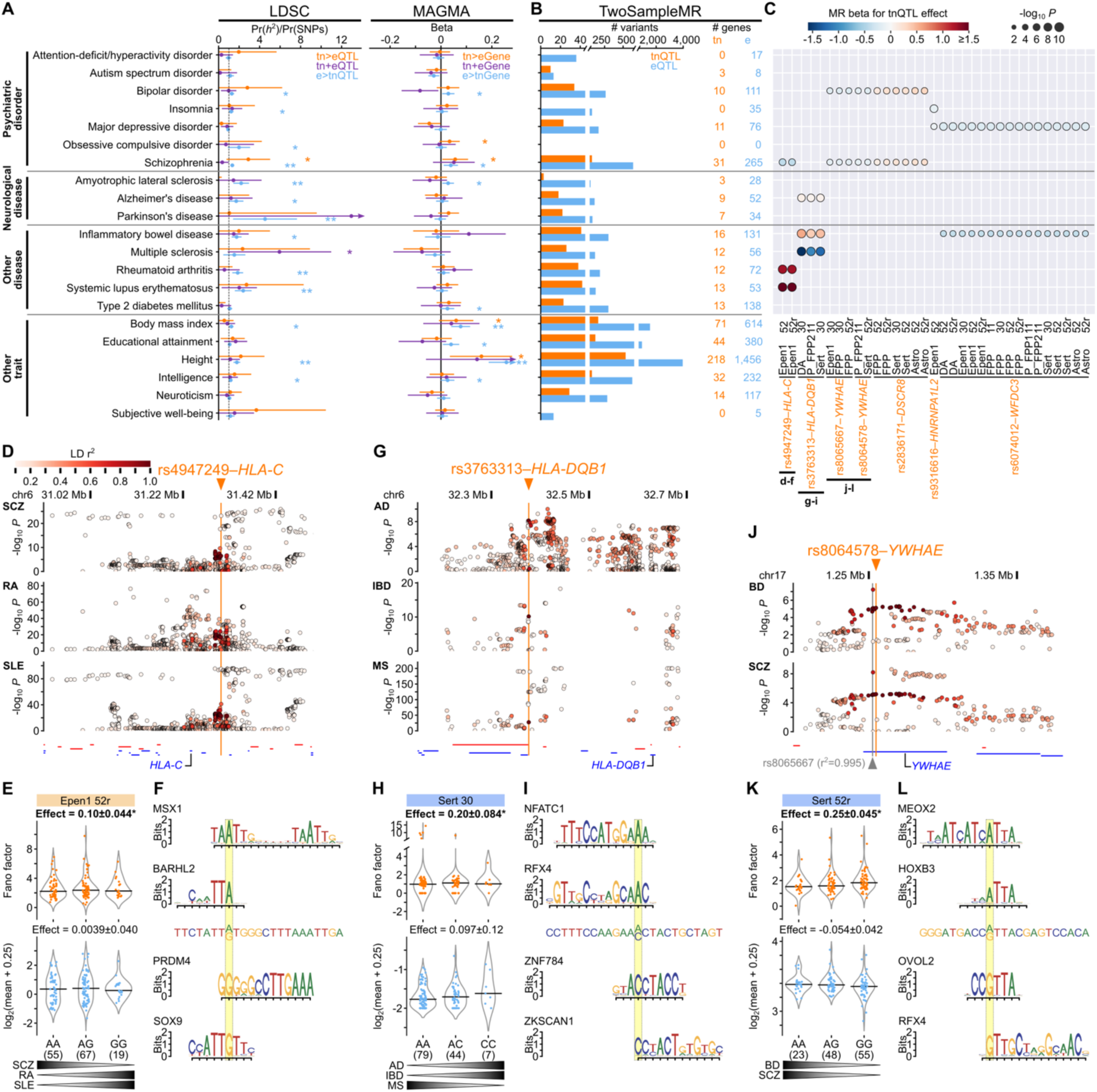
Enrichment in loci associated with various human complex traits and Mendelian randomization. (A) Results of the LDSC analysis testing for heritability enrichment in tn>eQTLs (orange), tn+eQTLs (purple), and e>tnQTLs (cyan) (left) and the MAGMA analysis evaluating enrichment of aggregated genetic association signals in tn>eGenes, tn+eGenes, and e>tnGenes (right) for various human complex traits. \**P* < 0.05; \*\**P* < 0.05/63 (i.e., significant after Bonferroni correction). The error bars indicate 95% confidence intervals. (B) The numbers of tnQTL and eQTL SNPs (left) and tnGenes and eGenes (right) whose causal association was indicated by a Mendelian randomization (MR) analysis. (C) The tn>eQTL SNP-tnGene pairs that showed causal association with multiple traits. The cellular conditions in which significant causal associations were observed are displayed. The colors and sizes of the circles indicate effect sizes in the MR analysis and -log_10_ *P* values for trait association, respectively. The corresponding panels of local plots are indicated below the pairs. (D-L) Local plots of tn>eQTL SNP-tnGene pairs that showed causal association with multiple traits. (D, G and J) show local Manhattan plots. Each circle indicates a SNP and is color-coded by LD with the lead tn>eQTL SNP (orange arrowhead) in the respective locus. Genes in each locus are shown at the bottom of the plot with the label of causal tnGene. Genes in red and blue indicate those coded on the forward and reverse strands, respectively. SNPs with *r*^2^ < 0.1 are not shown. (E, H, and K) are violin plots of the Fano factors (top) and mean expression levels (bottom) of the gene targeted by the lead tn>eQTL SNP in each locus per genotype. The posterior mean of estimated effect (slope) ± the standard deviation for each SNP is shown above the plot, and significant effects are in bold and marked with an asterisk. The horizontal black lines indicate the median. (F, I and L) indicate up to two TF consensus motifs potentially disrupted (top) or created (bottom) by the variant allele (right in the violin plots) of the lead tn>eQTL SNP. The SNP site and the surrounding sequences are shown between the TF consensus motif logos. AD, Alzheimer’s disease; BD, bipolar disorder; IBD, inflammatory bowel disease; MS, multiple sclerosis; RA, rheumatoid arthritis; SCZ, schizophrenia; SLE, systemic lupus erythematosus.

In the LDSC analysis, we observed significant enrichment of heritability for height, Parkinson’s disease, systemic lupus erythematosus (SLE), amyotrophic lateral sclerosis (ALS), rheumatoid arthritis (RA), and schizophrenia in e>tnQTLs after Bonferroni correction for multiple testing (**Figure 5A**, left, 21 traits x 3 QTLs = 63 tests in total). It would be reasonable to observe significant enrichment with a large effect size for Parkinson’s disease, where the importance of midbrain neurons in its pathology is well established. Likely reflecting their much smaller numbers than e>tnQTLs, there was no significant enrichment in tn>eQTLs and tn+eQTLs after multiple testing correction. Meanwhile, nominal enrichment (uncorrected *P* < 0.05) of schizophrenia heritability in tn>eQTLs and multiple sclerosis (MS) heritability in tn+eQTLs were observed. In the MAGMA analysis, e>tnGenes were enriched for the association signals for height and body mass index (BMI) after Bonferroni correction, whereas no significant enrichment was observed in tn>eGenes and tn+eGenes for any traits (**Figure 5A**, right). Nominal enrichment in tn>eGenes was found in height, schizophrenia, BMI, and obsessive-compulsive disorder (OCD). Overall, the numbers of tn>eQTLs and tn+eQTLs and their target genes seemed insufficient to identify robust enrichment. Nevertheless, for schizophrenia, SLE, and subjective well-being, we observed positive enrichment effect sizes that were consistently larger in tn>eQTLs and their targets than the other types of QTLs in both LDSC and MAGMA analyses (**Supplemental Table S4**).

### Exploration of tnQTLs causally associated with traits by Mendelian randomization

In addition to the above analysis examining the overall enrichment of trait association signals in the loci controlling transcriptional noise and their targets, we sought to identify transcriptional noise causally associated with traits by performing a Mendelian randomization analysis using TwoSampleMR^42^. By using tnQTL or eQTL SNPs as instrumental variables, we identified genes whose transcriptional noise or expression abundance is causally associated with the outcome phenotypes (**Figure 5B** and **Supplemental Table S5**). Although fewer than eGenes, we identified certain numbers of tnGenes shown to have causal effects for many traits.

Among these causal tnQTL SNP-tnGene pairs, a total of seven were found to be associated with the risks of multiple traits and involved tn>eQTLs (**Figure 5C** and local plots in **Figures 5D-L** and **Supplemental Figure S10**). Of these, two were found in the human leukocyte antigen (HLA) region, one being the magnitude of *HLA-C* transcriptional noise in day 52 Epen1 cells associated with the rs4947249 genotypes. In this locus, a larger noise of *HLA-C* was linked to a lower risk of schizophrenia and higher risks of SLE and RA (**Figures 5D-E**). The variant allele of this SNP was predicted to disrupt MSX1 and BARHL2 consensus motifs while creating potential binding sites of PRDM4 and SOX9 (**Figure 5F**). Since it is well known that the haplotypes of *C4* in the HLA region are associated with schizophrenia and autoimmune disease risks in opposite directions^43^, we examined the relationship between this SNP and the *C4* haplogroups and found that there is a link between them (**Supplemental Figure S11**). Therefore, the causal effect of this SNP would be mediated by *C4*, while tnQTL effects on *HLA-C* may also explain some of the risk mechanisms (detailed in **Discussion**). Another SNP in the HLA region, rs3763313, is associated with transcriptional noise of *HLA-DQB1*, whose larger noise was associated with higher Alzheimer’s disease and inflammatory bowel disease (IBD) risks and a lower MS risk (**Figures 5C**, **G**, and **H**). Thus, different directions of causal associations within autoimmune diseases were observed for this tn>eQTL-tnGene pair.

The motifs predicted to be disrupted or created by this SNP included those of NFACT1, RFX4, ZNF784, and ZKSCAN1 (**Figure 5I**). Four tn>eQTL-tnGene pairs in three loci (rs8065667-*YWHAE*, rs8064578-*YWHAE*, rs2836171-*DSCR8*, and rs9316616-*HNRNPA1L2*) showed causal associations with neuropsychiatric phenotypes that are known to be genetically correlated^44^ (**Figures 5C** and **J-L**, and **Supplemental Figures S10A**-**F**). The rs8065667 and rs8064578 SNPs in the *YWHAE* locus were in strong linkage disequilibrium with each other (*r*^2^ = 0.995), while either was identified as the lead SNP in different cellular conditions (**Figure 5C**), and smaller noise of *YWHAE* was associated with risks of schizophrenia and bipolar disorder (**Figures 5J** and **K**). Causal associations common to schizophrenia and bipolar disorder were also observed for the rs2836171-*DSCR8* pair on another chromosome. Contrary to *YWHAE*, larger *DSCR8* noise was associated with the risks of these disorders (**Supplemental Figures S10A** and **B**). The rs9316616-*HNRNPA1L2* pair was associated with another group of genetically correlated neuropsychiatric phenotypes, major depressive disorder (MDD) and insomnia, where smaller noise of this gene increases their risks (**Supplemental Figures S10D** and **E**). Unlike these tn>eQTL SNP-tnGene pairs that were significant only in specific cellular conditions, the rs6074012-*WFDC3* pair showed causal associations with MDD and IBD in almost all types of cells (**Figures 5C**, and **Supplemental Figures S10G**, and **H**). Representative TF consensus motifs potentially disrupted or created by these SNPs (rs8064578, rs2836171, rs9316616, and rs6074012) are displayed in **Figure 5L**, and **Supplemental Figure S10C**, **F**, and **I**.

### Transcriptional noise dysregulation in schizophrenia patient and model brains

Following our analysis of the roles of tnQTLs in various human complex traits, we further investigated possible dysregulation of transcriptional noise in the disease-responsible organ, focusing on schizophrenia, for which LDSC and MAGMA analyses consistently showed nominal enrichment in tn>eQTLs and tn>eGenes with effect sizes larger than those of e>tnQTLs/e>tnGenes (**Figure 5A**), specific causal tn>eQTL associations were identified (**Figures 5B** and **C**), and single-nucleus(sn)/scRNA-seq datasets for human case-control samples and disease models were available. To this end, we analyzed data from the study by Batiuk et al.^45^, which analyzed postmortem human brain neuronal nuclei from the Brodmann area 9 of the dorsolateral prefrontal cortex (DLPFC, **Figure 6A**; 79,403 cells from 9 schizophrenia cases and 117,819 cells from 14 controls), and the study by Chen et al.^46^, which analyzed prefrontal cortex (PFC) and striatum cells of mice with heterozygous knockout of a histone modifier gene *Setd1a*, whose human ortholog *SETD1A* is associated with schizophrenia with robust statistical significance and a large effect size^47–49^ (**Figure 6B**, 3,557 PFC and 6,290 striatum cells from two *Setd1a*^+/-^ and 5,145 PFC and 8,400 striatum cells from two wildtype mice). Comprehensive identification of genes that differ in transcriptional noise between cases and controls or mutant and wildtype mice, i.e., differentially noisy genes (DNGs), and differentially expressed genes (DEGs) with significant mean differences was performed by using the Bayesian Analysis of Single-Cell Sequencing data (BASiCS) package^50^ (see **STAR Methods** for details). In this analysis, significant DNGs were identified by calculating residual over-dispersion, a measure of the excess cell-to-cell variation in a cell population not confounded by mean expression, instead of the Fano factor, a metric of transcriptional noise of each gene in an individual used in the analysis of tnQTLs.

**Figure 6.**
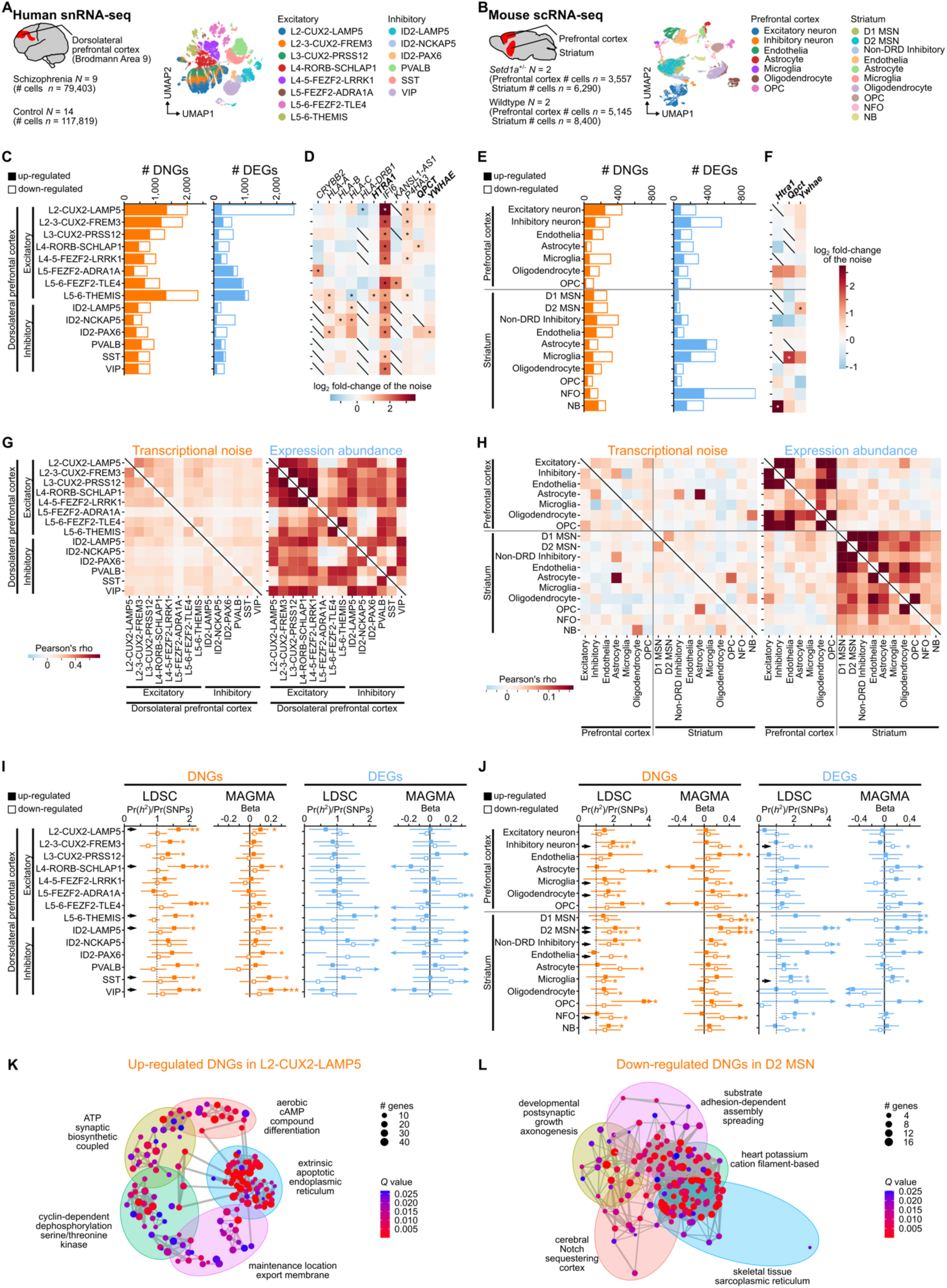
Transcriptional noise dysregulation in schizophrenia patient and model brains. (A and B) Datasets used. We used human snRNA-seq (A) and mouse scRNA-seq (B) data from the indicated numbers of cells and individuals. The uniform manifold approximation and projection (UMAP) representations of the single-nucleus and single-cell data and cell cluster labels are shown in the right parts of the panels. (C-F) Differentially noisy genes (DNGs) and differentially expressed genes (DEGs) in schizophrenia patient and model brains. The bar plots show the numbers of up-regulated (colored) and down-regulated (white) DNGs and DEGs in each cell type in the human postmortem DLPFC (C) and mouse PFC and striatum (E). (D) and (F) respectively indicate DNGs in humans (D) and mice (F) for which causal associations were detected in our Mendelian randomization analysis. The log_2_ fold changes of the noise in each cell type are shown as heatmaps. DNGs identified in both humans and mice are in bold. Significant DNGs in the BASiCS analysis are marked with asterisks. The black diagonal lines indicate genes not expressed in the corresponding cell type. (G and H) Correlation heatmaps of case-control log_2_ fold changes in transcriptional noise or expression abundance across cell types in human (G) and mouse brains (H). In this analysis, data of all genes, not restricted to DNGs or DEGs, were used. (I and J) Enrichment analysis of schizophrenia genetic association signals in DNGs and DEGs by LDSC and MAGMA. We separately analyzed up- and down-regulated genes in each cell type in human (I) and mouse brains (J). Enrichment effect sizes and the 95% confidence intervals are shown. The DNG and DEG sets that showed enrichment (uncorrected *P* < 0.05) in both LDSC and MAGMA are indicated with black arrows. \**P* < 0.05; \*\**P* < 0.05/number of cell types tested (14 in (I) and 17 in (J)). (K and L) Network visualization of GO terms enriched in the up-regulated DNGs in human DLPFC L2-CUX2-LAMP5 neurons (K) and the down-regulated DNGs in mouse striatal D2 MSN (L). Node sizes and colors indicate the numbers of hit genes and *Q* values, respectively.

Based on the cell cluster labels in the original publications^45,46^, we comprehensively detected DNGs and DEGs in each of the 14 cell-type clusters in the human postmortem brain data and the 17 cell types (seven in PFC and ten in striatum) in the mouse model data (**Figures 6A** and **B**). In the analysis of human postmortem brain data, 556-2,335 DNGs were found in each cell type (**Figure 6C** and **Supplemental Table S6**). The largest numbers of DNGs were observed in L5-6-THEMIS (*THEMIS*-expressing layer 5-6 excitatory neurons), followed by L2-CUX2-LAMP5 (*CUX2*- and *LAMP5*-expressing layer 2 excitatory neurons) and L2-3-CUX2-FREM3 (*CUX2*- and *FREM3*-expressing layer 2-3 excitatory neurons) (**Figure 6C**, left). Overall, more DNGs were found in excitatory neurons than in inhibitory neurons. There were no remarkable disparities in the numbers of genes exhibiting up- and down-regulated transcriptional noise in schizophrenia brains. For DEGs, we again found that their numbers were largest in the superficial (L2-CUX2-LAMP5) and deep layer (L5-6-THEMIS and L5-6-FEZF2-TLE4: *FEZF2*- and *TLE4*-expressing layer 5-6) excitatory neurons. In contrast to DNGs, marked biases toward up- or down-regulated DEGs were observed in these cell types (**Figure 6C**, right). Of the 31 tnGenes whose causal association with schizophrenia was shown in the Mendelian randomization analysis (**Figure 5B**), 20 were found to be expressed in this dataset, of which 11 genes were found as DNGs in one or more postmortem brain cell types (**Figure 6D**). Many of these DNGs showed cell type preference, while significant up-regulation of *IFI6* transcriptional noise was common except for L5-FEZF2-ADRA1A (*FEZF2*- and *ADRA1A*-expressing layer 5 excitatory neurons) and PVALB (parvalbumin-expressing inhibitory neurons), and up-regulated noise of *P4HA3* was found in many types of excitatory neurons. In the brains of *Setd1a* deficient mouse model, as in human schizophrenia, the number of DNGs was largest in excitatory neurons in the PFC. In the striatum, DNGs were most abundant in inhibitory neurons not expressing dopamine receptors D1 and D2 (non-DRD inhibitory, **Figure 6E**, left). Unlike DNGs, the largest numbers of DEGs in the mutants were observed in inhibitory neurons in the PFC and newly formed oligodendrocytes (NFO) in the striatum (**Figure 6E**, right). Among 12 of the 28 putative causal tnGenes whose orthologs are expressed in the mouse brain, cell-type specific up-regulation of transcriptional noise in three genes, *Hitra1*, *Qpct*, *Ywhae*, were observed in the schizophrenia model (**Figure 6F**). All of these three genes were also identified as up-regulated DNGs in one or more DLPFC cell types in human schizophrenia (indicated in bold in **Figure 6D**), while significant up-regulation was observed only in the striatal cells in the mouse model.

Next, we analyzed the correlations of case-control fold changes in transcriptional noise or expression levels between each cell type pair to understand whether differences between cases and controls are shared or not shared across cell types. We found a general trend that the fold changes in expression abundance are correlated across cell types within a brain region (**Figure 6G**, right, and **Figure 6H**, right), whereas the noise differences are less correlated with few exceptions such as a strong correlation between mouse PFC and striatum astrocytes (**Figure 6G**, left, and **Figure 6H**, left). These contrasting results would suggest that dysregulation of transcriptional noise in schizophrenia is more cell type-specific than that of expression abundance and thus may be more informative in specifying disease-relevant cell types.

In addition to the analyses based simply on the numbers of DNGs and DEGs shown in **Figures 6C** and **E**, we also evaluated if DNGs and DEGs in each cell type were enriched for association signals in the schizophrenia GWAS^51^. As in the analysis of tnQTLs, we used LDSC^39,40^ and MAGMA^41^, and the up- and down-regulated genes were analyzed separately. Enrichment of schizophrenia association signals at uncorrected *P* < 0.05 in the human postmortem brain dataset was commonly detected by LDSC and MAGMA in the DNGs up-regulated in the following six cell types (**Figure 6I**, left, indicated by the black arrows; **Supplemental Table S7**): L2-CUX2-LAMP5; L4-RORB-SCHLAP1 (*RORB* and *SCHLAP1*-expressing layer 4 excitatory neurons); L5-6-THEMIS; ID2-LAMP5 (*ID2* and *LAMP5*-expressing inhibitory neurons); SST (somatostatin-expressing inhibitory neurons); VIP (vasopressin-expressing inhibitory neurons). On the other hand, neither the down-regulated DNG nor both directions of DEG sets showed consistent enrichment in the LDSC and MAGMA analyses (**Figure 6I**, right). In the *Setd1a*^+/-^ mouse brain data, enrichment of schizophrenia association signals both in the LDSC and MAGMA analyses was preferentially observed in down-regulated DNGs, which include inhibitory neurons, microglia, and oligodendrocyte in the PFC and D2 MSN (*Drd2*-expressing medium spiny neurons), non-DRD inhibitory neurons, endothelia, and NFO in the striatum (**Figure 6J**, left). In the striatal D2 MSN, up-regulated DNGs were also enriched for schizophrenia association signals. As in human postmortem brains, fewer DEG sets showed consistent enrichment in the *Setd1a*^+/-^ mouse brains than DNGs, where only two down-regulated DEG sets of PFC inhibitory neurons and striatal microglia were supported by both LDSC and MAGMA (**Figure 6J**, right). Overall, enrichment of schizophrenia association signals was observed more in the DNG sets, which include those in the superficial and deep layer cortical excitatory neurons and D2 MSN implicated in schizophrenia pathology or therapeutic targets^45,52–58^, than in the sets of DEGs (14 DNG sets vs. 2 DEG sets).

When we performed a GO enrichment analysis using clusterProfiler^24^ to evaluate the properties of up-regulated DNGs in human L2-CUX2-LAMP5 and down-regulated DNGs in mouse D2 MSN, which showed consistent and significant enrichment in the above analyses, we found that these DNG sets were both enriched for terms related to neuronal components, such as synapses and axons (**Figures 6K** and **L**). A more specific enrichment analysis focusing on synaptic communications using Synaptic Gene Ontology (SynGO^59^) terms revealed that presynaptic and postsynaptic terms were particularly enriched in up-regulated DNGs in L2-CUX2-LAMP5 and down-regulated DNGs in D2 MSN, respectively (**Supplemental Figure S12**). On the other hand, significant enrichment of synaptic terms was observed neither in the opposite directions of DNGs nor any DEGs in these cell types. These results, showing the enrichment of pathways and components whose involvement in the schizophrenia pathophysiology has been established, could further support the role of transcriptional noise dysregulation in this disorder.

## Discussion

In this study, we took advantage of large-scale scRNA-seq data from iPSC-derived brain cells and comprehensively mapped tnQTLs, the genomic regions regulating cell-to-cell variation in gene expression abundance. Although a similar previous attempt^60^ analyzing scRNA-seq data of 5,597 cells from 53 individuals profiling 9,957 protein-coding genes failed to identify statistically robust tnQTLs at FDR < 0.1, we analyzed more individuals (*N* = 155), cells (total *n* = 795,661), and genes (total *n* = 16,954) by utilizing mash^26^, a recently developed statistical method that can integrate scRNA-seq data from multiple conditions, and identified a total of 101,024 tnQTL-gene pairs at the level of local false sign rate (lfsr)^27^ < 0.05. This would be consistent with the simulated results of QTL detection using scRNA-seq data by powerEQTL^61^, which showed a substantial difference in statistical power between the above two conditions (**Supplemental Figure S13**).

In parallel to the tnQTL detection, we identified eQTLs genome-wide using an equivalent pipeline. Our comparison of tnQTLs and eQTLs showed that the majority of tnQTLs have significant eQTL effects (**Figure 2E**). This may be reasonable given the assumed mechanism through which QTLs modulate transcriptional noise. For example, an increase in transcriptional noise could be explained by a larger amplitude of transcriptional bursts, and in this case, the average expression level should inevitably increase unless the duration and/or frequency of the bursts decreases. Thus, the tn+eQTLs we identified in this study may have such an effect on the amplitude, whereas tn>eQTLs, which were fewer in our analysis, could confer fine-tuning of burst amplitudes, durations, and frequencies in a way that does not significantly change the mean. This speculation can be interpreted together with recent literature and the results of our enrichment analysis. A study of transcriptional burst kinetics reported the importance of both promoters and enhancers, where the former is more associated with burst sizes and the latter with burst frequencies^62,63^. In our enrichment analysis of functional genomic elements, tn+eQTLs were most prominently enriched in many regulatory elements including promoters and enhancers, while tn>eQTLs are not significantly enriched in TATA-containing promoters (**Figure 4A**). Therefore, if SNPs in these promoter regions specifically affect burst sizes, our observation of no significant tn>eQTL enrichment might be reasonably explained.

The different characteristics of tn>eQTLs compared to the other types of QTLs (tn+eQTLs and e> tnQTLs) were also indicated in other analyses, including those of TF-binding site enrichment (**Figures 4C-H**), population frequencies (**Supplemental Figure S9A**), evolutionary conservation (**Supplemental Figure S9B**), LoF-constraint of target genes (**Supplemental Figure S9D**), and correlations across cellular conditions (**Figures 3A** and **B**). In addition, our analysis of DNGs and DEGs in schizophrenia patient and model brains showed that DNGs are less shared across cell types than DEGs (**Figures 6G** and **H**), supporting that transcriptional noise and expression abundance undergo different regulations with the former being more cell type-specific. It was also shown that DNG and DEG differ in the number of differential genes and in the enrichment of genetic association signals for schizophrenia, where the cell types indicated in the DNG analysis included superficial and deep layer excitatory neurons, specific types of inhibitory neurons, and D2 MSNs (**Figures 6C, E, I**, and **J**). These are generally consistent with the cell types whose involvement in the schizophrenia pathology was supported by both large-scale common SNP GWAS and rare protein-coding variant sequencing study for schizophrenia^47,64^. Therefore, analyses focusing not only on expression levels but also on transcriptional noise might enhance our understanding of cell types associated with the pathophysiology of various human diseases, not restricted to schizophrenia. In the context of the association between transcriptional noise and human diseases and traits, we also performed LDSC and MAGMA analyses of tn>eQTLs, tn+eQTLs, and e>tnQTLs and their target genes (**Figure 5A**). These analyses showed enrichment with particularly large enrichment fold changes in tn>eQTLs for several traits, such as schizophrenia, SLE, and subjective well-being, tough the statistical significance was modest and did not achieve the threshold after multiple testing correction. Subsequent Mendelian randomization analysis identified a number of tnQTL SNP-tnGene pairs with causal effects (**Figure 5B**). Of these, we identified six independent pairs associated with multiple traits (**Figure 5C**), which included the rs4947249-*HLA-C* pair linked to schizophrenia and autoimmune disease risks in opposite directions. In addition to the well-known roles of HLA genes in the immune system, human HLA class I genes, consisting of *HLA-A*, *HLA-B*, and *HLA-C*, are expressed in the developing brain^65,66^ and adult dopaminergic neurons^67^ and have been implicated in various biological processes such as cellular differentiation, synapse elimination, and reactive cell death^67–69^. On the other hand, it should be recognized that the disease association signals in the HLA region, which is comprised of complex linkage disequilibrium blocks, can be mediated by effects other than those on the transcriptional noise of the *HLA-C* gene observed in this study, such as the copy number and gene expression abundance of the *C4* genes, whose haplogroups are shown to be associated with this rs4947249 polymorphism (**Supplemental Figure S11**). Nevertheless, given that the involvement of monoaminergic neurons in schizophrenia has been consistently supported by pharmacological and neural circuit studies^70^, our observations raise the possibility that a part of the causal effect of genetic variants in this locus on schizophrenia risk may be explained by their impact on the transcriptional noise of *HLA-C* in these neurons. Also, while we were unable to detect *C4* expression in the cells analyzed in this study, nor were we able to dissect *C4A* and *C4B* in the analysis of scRNA-seq data, it would be beneficial to evaluate the effects of the associated SNP(s) on the transcriptional noise of *C4* genes in other cell types.

Considering the limitations of this study, first, we only analyzed specific types of cells, and a broader range of cell types and tissues need to be analyzed to more comprehensively understand the regulation of transcriptional noise by tnQTLs. For a specific example, as mentioned above, this study identified a tnQTL in the HLA locus encompassing the *C4* genes, which was suggested to be causally associated with diseases; however, the *C4* genes were not expressed in the cells analyzed in this study. Therefore, we were unable to assess whether this region is associated with *C4* transcriptional noise and, if so, whether it could explain a part of the causal relationship. Further analyses of other cell types, which are within the scope of our future studies, would provide insights into this unsolved question. Second, although our study was able to identify a number of tnQTLs by analyzing a total of 795,661 cells derived from 155 donors, it is expected that more loci could be identified by increasing the sample size. In particular, since our results indicate that many tnQTLs also have properties as eQTLs, a larger study is needed to more clearly delineate the role of QTLs that show effects exclusively on transcriptional noise. Third, our study only analyzed SNPs that are found at a certain frequency in the population and did not characterize possible roles of rare variants, which may have larger effect sizes. It is also possible that rare somatic variants that exist only in a subset of cells shape the transcriptional noise. Analyses of these types of rare variants will provide a more comprehensive picture of the genetic loci controlling transcriptional noise.

Nevertheless, this study, which identified tnQTLs on a large scale by leveraging the information from scRNA-seq data, provides a catalog of genetic variants controlling a molecular phenotype that has not been thoroughly investigated in previous QTL studies. Furthermore, our characterization of tnQTLs by integrating other types of genomic datasets reveals relationships between tnQTLs and eQTLs, as well as enrichment patterns of QTLs having both or either of these effects in regulatory elements and human complex trait-associated loci. By advancing research aiming to fully extract information from big omics data by effectively combining different layers of datasets and analyzing them from unconventional viewpoints, we will be able to obtain deeper insights into the mechanisms of various biological phenomena, such as gene expression regulation and disease pathogenesis.

## Acknowledgements

We thank Daisuke Yamaguchi and Hao Zhang from BITS Co., Ltd., for their assistance in the construction of the analytical environment and Professor Kazuki Nakao at Institute of Experimental Animal Sciences, Osaka University, for his insightful advice. We made use of data produced by the HipSci Consortium funded by the Wellcome Trust and the MRC. This research was supported by the RIKEN Center for Brain Science, RIKEN Student Researcher Program (Y.N.), and the following funding agencies: AMED under grant numbers JP23km0405214 (A.T.), JP24tm0524001 (A.T.), and JP24wm0625001 (A.T.); JSPS KAKENHI under grant numbers JP21H02855 (A.T.), JP22K15785 (N.H.), JP22K20751 (S.M.) and JP24K18729 (S.M.). Computations in this study were performed on the NIG supercomputer at ROIS National Institute of Genetics.

## Author contributions

N.H. and A.T. designed the study and wrote the manuscript. N.H., S.M., Y.N., T.H., H.Y., E.K., and J.U. contributed to the data acquisition, analyses, and manuscript refinement. T.T. provided conceptual advice. A.T. supervised the entire study. All the authors read and approved the manuscript.

## STAR Methods

### Ethical statement

The study was approved by the Wako first ethics committee of RIKEN (approval number: Wako1 2021-001(6)) and conforms to the provisions of the Declaration of Helsinki. We used human genotype and transcriptomic data publicly available without access restrictions detailed below.

### Acquisition of iPSC-derived midbrain cell data

The scRNA-seq data (a count matrix) of 1,027,398 iPSC-derived midbrain cells from 215 individuals and the genotyping data of 156 individuals were downloaded from the European Nucleotide Archive (ENA) with the accession numbers summarized in **Supplemental Table S8**. This dataset was generated through the Human Induced Pluripotent Stem Cells Initiative (HipSci) project. The detailed methods for the establishment of iPSC lines and differentiation into midbrain cells are described in Ref 22. Briefly, after the collection of primary fibroblasts from skin biopsies from healthy European volunteers who had consented to the research usage, iPSCs were established using the Sendai reprogramming kit or episomal plasmids, and then quality controlled for the following items: level of pluripotency, number of copy number abnormalities, and the ability of differentiation to each of the three germ layers. Neuronal differentiation of the pooled iPSC lines to a midbrain lineage was performed according to the protocol found at https://www.protocols.io/view/generation-of-ipsc-derived-dopaminergic-neurons-bjpgkmjw.

### Genotyping data processing

The raw SNP genotyping data for each of the 156 individuals called by the HumanCoreExome BeadChip (Illumina, USA) are available in the ENA in the VCF format. We processed this dataset by following the procedures described in the GTEx project^71^ using bcftools (version 1.9), vcftools (version 0.1.15), and plink (version 1.90b). After confirming the agreement between the gender information in the metadata and the sex inferred from the genotypes, we first merged individual donors’ VCF files using bcftools *merge*. There were a total of 536,047 genotyped sites in the merged dataset. We then performed initial quality controls using plink with the following criteria: call rate ≥ 95% (*n* of the remaining sites = 503,953), polymorphic (*n* = 307,262), and *P* values for deviation from Hardy-Weinberg equilibrium ≥ 1 × 10^-6^ (*n* = 304,497). Since Y chromosome genotype data were not available for some male donors, we did not include the Y chromosome in our analyses. All X chromosome variants analyzed were in the non-pseudoautosomal region. Using the quality-controlled genotyping dataset, we performed haplotype phasing of each autosomal and X chromosome by SHAPEIT^72^ (version 2.r837). The *--effective-size* parameter of SHAPEIT was set to 11418 according to the developer’s instruction. The X chromosome haplotypes were phased with the *--chrX* option. Heterozygous X chromosome genotypes of male donors were masked and considered as missing alleles. Based on this phased dataset, we performed an imputation of ungenotyped variants using IMPUTE2^73^ (version 2.3.2) [https://doi.org/10.1371/journal.pgen.1000529]. The chunk ± buffer region sizes were set to 3 Mb ± 250 kb for autosomes and 5 Mb ± 300 kb for X chromosome. The *-Ne* parameter was set to 20000, according to the developer’s guide. Since the reported ethnicity of all donors was European, the European reference panel of the 1000 Genomes Project Phase 3 was used (see the URL section for file accession) for the phasing and imputation. There were a total of 25,216,663 variants in the genotyped + imputed dataset. We then applied the following filtering using bcftools (version 1.9): IMPUTE2 info score ≥ 0.4 (*n* of the remaining variants = 13,135,257), length of insertions and deletions ≤ 50 bp (*n* = 13,134,196), minor allele frequency ≥ 0.01 (*n* = 9,540,431), and Hardy-Weinberg equilibrium deviation *P* ≥ 1 × 10^-6^ (*n* = 9,491,861). We annotated the set of 9,491,861 SNPs with reference SNP IDs (rsIDs) using bcftools *annotate* (version 1.9) based on dbSNP151 (http://hgdownload.soe.ucsc.edu/goldenPath/hg19/database/snp151.txt.gz) and used this dataset for the downstream analyses. We confirmed that all donors included in this study were of European ancestry by performing a principal component analysis of the studied subjects together with the European, East Asian, and African superpopulations of the 1000 Genomes Project Phase 3 data (build 37, 2504 samples; https://www.cog-genomics.org/plink/2.0/resources#phase3_1kg) using plink (versions 2.00a5.10 and 1.90b) and eigensoft (version 8.0.0) (**Supplemental Figure S3**).

### Single-cell RNA-seq count matrix processing and transcriptional noise quantification

The scRNA-seq data of midbrain cells differentiated from human iPSCs^22^ were downloaded from https://zenodo.org/record/4651413 (day11.h5, day30.h5, and day52.h5 of the Version v3). From the downloaded HDF5 files, we retrieved the raw unique molecular identifier (UMI) count matrix and cell type annotations using scanpy (version 1.8.1). We then split the matrix into data for each of the cell types determined in the Jerber et al. study. After excluding cells that failed cell type labeling and rare cell types with small numbers of cells, the following datasets for 17 cellular conditions were used in this study: days 30 and 52 dopaminergic neurons (DA); days 30 and 52 ependymal-like cells (Epen1); days 11, 30 and 52 floor plate projenitors (FPP); day 11 proliferating FPP type 1 and 2 (P_FPP1 and P_FPP2); days 30 and 52 serotonergic-like neurons (Sert); day 52 astrocyte-like cells (Astro); day52 cells include those treated with rotenone to mimic oxidative stress conditions and untreated cells. Gene-level filtering was performed by excluding those expressed in < 1% of cells in each cellular condition, as in the study by Jerber et al. We then subjected the dataset for each condition to an adjustment for the dependency of the mean and variability in gene expression using the *BASiCS_MCMC()* function of the BASiCS^50^ package (version 2.6.0) with the following parameters: *WithSpikes = F, N = 20000, Thin = 20, Burn = 10000, Regression = T*, and the pool IDs as BatchInfo. The adjusted (denoised) gene expression counts in each cellular condition were then calculated using *BASiCS_DenoisedCounts()*. Successful denoising was confirmed by comparing over-dispersion (*δ*) and residual over-dispersion (*ε*) (**Supplemental Figure S1**). These preprocessing procedures were performed by using the data from 215 donors, including those without genotyping data. Using the denoised count matrices for the 17 conditions created as described above, we quantified the transcriptional noise of each gene in each donor by calculating the Fano factor (variance/mean). This means that we applied a conservative double correction for potential confounding by mean, by performing a mean-variability adjustment with BASiCS and calculating the Fano factor, in which the variance is divided by the mean. When a gene is expressed in none of all cells of a given donor and therefore the expression mean was 0, the corresponding Fano factor was set to 0.

### GSEA of genes with large and small transcriptional noise

We performed Gene Set Enrichment Analysis (GSEA) using the R package clusterProfiler^24^ (version 4.2.2) based on the mean transcriptional noise levels (Fano factors) of each gene across the cohort of 215 donors (including individuals with no genotyping data) to characterize biological features of genes with large and small transcriptional noise. We subjected the undifferentiated day11 P_FPP1 and day11 P_FPP2 and the differentiated day52 DA and day52 Sert datasets for this analysis. We used the *gseGO*() function of clusterProfiler^24^ with the org.Hs.eg.db (version 3.14.0) annotation using the following parameters after generating random number seed by *set.seed*(1): *ont* = “ALL”, *OrgDb* = org.Hs.eg.db, *exponent* = 1, *minGSSize* = 10, *maxGSSize* = 500, *eps* = 0, *pvalueCutoff* = 0.05, *pAdjustMethod* = “BH”, and *seed* = T. The dot plots of the GO terms enriched in noisy or stable genes were produced by using the R packages enrichplot (version 1.14.2) and ggplot2 (version 3.4.4). For visualization, we also used the seaborn (version 0.11.0) and matplotlib.pyplot (version 3.3.2) modules of python (version 3.8.5).

### Genome-wide identification of tnQTLs and eQTLs

#### Analysis of individual datasets by FastQTL

In each of the individual datasets for the 17 cellular conditions, donors for whom data of 5 or more single cells were available were used for the QTL detection. One donor (HPSI1113i-uofv_1) was excluded from the subsequent analyses as this individual did not meet this criterion in all 17 conditions. Consequently, a total of 155 donors were included in the QTL analysis. We first performed genome-wide identification of tnQTLs and eQTLs in each condition using FastQTL^25^ (enhanced version of 2.184; https://github.com/francois-a/fastqtl), following the procedures used in the GTEx project^71^. The gene × donor matrices for transcriptional noise (Fano factor) or expression abundance (log_2_(mean + 0.25)) were used as the input for quantitative traits. Genic regions were defined using the GENCODEv38lift37 comprehensive annotation (https://ftp.ebi.ac.uk/pub/databases/gencode/Gencode_human/release_38/GRCh37_ mapping/gencode.v38lift37.annotation.gtf.gz). We used this version because the VCF files downloaded from ENA were based on the GRCh37 coordinates. When multiple entries were assigned to a single gene, we selected the one with the largest start-to-end distance. Genes in the Y chromosome and mitochondrial genome were not included in our analysis. The input genotype dataset was prepared by calculating the alternate allele dosage (DS) for each SNP in each donor using the estimated posterior genotype probabilities (GP) with the equation DS = P(0/1)+2*P(1/1). For the covariates, we used the first three principal components of the genotypes, the genotyping platforms, the sex information, and the first 15 probabilistic estimation of expression residuals (PEER) factors of the Fano factors for the tnQTL analysis and log_2_(mean + 0.25) for the eQTL analysis, which were calculated using the R package peer^74^ (version 1.0). After quantile normalization of the Fano factors and expression abundance (log_2_(mean + 0.25)) to the standard normal distribution, tnQTLs and eQTLs were identified for each of the 17 conditions using linear regression implemented in FastQTL^25^ with ±1 Mbp *cis*-windows. Following the GTEx protocol, we initially calculated nominal *P* values by running the “nominal pass” of FastQTL^25^ using *nominal* options. We then ran the “permutation pass” using *permutations* option to obtain empirical *P* values extrapolated from a Beta distribution fitted to adaptive permutations with the –permute 1000-10000 setting. Additional multiple testing correction across genes was performed by using the Storey & Tibshirani FDR procedure^75^.

#### Bayesian multi-condition analysis by mash

Multivariate adaptive shrinkage (mash^26^) is a Bayesian statistical approach that utilizes information shared across multiple conditions to improve the effect estimates and increase detection power. Since we analyzed 17 cellular conditions that are distinct but belong to the same lineage of neurons or their precursors, we thought that this framework could reasonably improve QTL detection capability. According to the developer’s instructions for mashr, the R package of mash^26^ (version 0.2.57, https://stephenslab.github.io/mashr/articles/eQTL_outline.html), we prepared the mash input files that contain the effect sizes (slope in the FastQTL output) and their standard errors for each tnQTL- or eQTL-gene pair in the 17 cellular conditions using individual FastQTL output files. In the FastQTL outputs, “NA” was assigned to the slopes and standard errors for non-expressed genes, and “NaN” was assigned to the standard errors for genes that were expressed but had a slope of 0. These missing effect sizes and standard errors were set to 0 and 1, respectively, according to a previous study^76^.

From these modified mash inputs, we randomly subsampled 50,000 variant-gene pairs as the “random set” under *set.seed(1)* and estimated the correlation structure in the null tests from this random set using *estimate_null_correlation_simple()*. We parallelly defined the most significant tnQTL or eQTL per gene as the “strong set”. Data-driven covariances in the strong set were calculated with their first five principal components using *cov_pca()* and *cov_ed()*. The canonical covariances in the random set were calculated using *cov_canonical()*. After fitting the mash model to the random set with the data-driven and the canonical covariances using *mash()*, the fitted model was applied to all variant-gene pairs by re-running *mash()* with the *g = get_fitted_g(m)* and *fixg = T* options, which specifies the learned model *m*. We separately analyzed each chromosome to reduce the computation burden. The posterior mean of effect estimate and the corresponding local false sign rate (lfsr), which is analogous to the local false discovery rate but measures confidence in the sign of each effect in the context of multiple hypothesis testing^27^, and posterior standard deviation were calculated using the *get_pm()*, *get_lfsr(),* and *get_psd()* functions. The posterior mean of effect estimate, lfsr, and posterior standard deviation for non-expressed genes were replaced with 0, 1, and 0, respectively. We considered QTL-gene pairs with lfsr < 0.05 in one or more of the 17 conditions to be significant and extracted them using *get_significant_results()*. In the integrated dataset of the 17 conditions, there were in total 101,024 and 1,224,228 significant tnQTL- and eQTL-gene pairs, respectively.

#### Stratification into tn>eQTLs, tn+eQTLs and e>tnQTLs

The significant tnQTL- and eQTLs-gene pairs in the integrated dataset of the 17 conditions detected by mash^26^ were stratified into tn>eQTLs, for which only the tnQTL effect was significant, tn+eQTLs, for which both the tnQTL and eQTL effects were significant, and e>tnQTLs, for which only the eQTL effect was significant. The distance between the TSS (5’ end) of the target gene and each tn>eQTL, tn+eQTL, or e>tnQTL SNP was calculated using a custom script based on the GENCODEv38lift37 annotation in a way aware of the transcription direction.

### Comparison of tnQTLs and eQTLs across the 17 conditions

Pairwise Pearson’s rhos of the posterior mean of tnQTL or eQTL effect estimates across the 17 cellular conditions were calculated by using the pandas (version 1.3.5) python module. For tn+eQTLs, we analyzed correlations of tnQTL and eQTL effects separately using the same dataset of tn+eQTL-gene pairs. In the comparison of rotenone-treated and untreated day 52 DA or day 52 Sert cells, we focused on tn>eQTLs targeting genes commonly expressed regardless of the rotenone treatment. GO enrichment analysis of the tn>eGenes promoted or suppressed by rotenone treatment in the day 52 DA or day 52 Sert cells was performed by using the *enrichGO()* function of clusterProfiler^24^ (version 4.2.2) with the following parameters: *OrgDb* = org.Hs.eg.db (version 3.14.0), *ont* = “ALL”, *pvalueCutoff* = 1, *pAdjustMethod* = “BH”, *qvalueCutoff* = 1, *minGSSize* = 10, and *maxGSSize* = 500.

### Enrichment analyses of tn>eQTLs, tn+eQTLs, and e>tnQTLs in functional genomic elements

We examined the enrichment of tn>eQTLs, tn+eQTLs, and e>tnQTLs in varisous functional genomic elements using GARFIELD^29^ (version 2). GARFIELD requires 1) a set of genome-wide association *P* values, 2) genomic coordinates for regulatory annotations of interest, 3) lists of linkage disequilibrium (LD) tags for each variant from a reference population, and 4) the distance of each variant to the nearest TSS. The sets of genome-wide association *P* values for tn>eQTLs and e>tnQTLs was prepared by selecting the smallest lfsr (significance statistics in mash^26^) for each SNP for the corresponding QTL effect when a SNP was associated with multiple genes. For tn+eQTL, similarly, we first selected the smallest lfsr for tnQTL effects and eQTL effects for each SNP when a SNP was associated with multiple genes. Then, for each SNP, the larger of these smallest tnQTL effect lfsr and smallest eQTL effect lfsr was considered the association significance for that SNP. In the tn+eQTL association *P* dataset produced by this procedure, all significant tn+eQTLs detected by mash^26^ had *P* values less than 0.05, whereas all others had *P* values greater than or equal to 0.05. Genomic coordinates for functional genomic annotations were obtained from various resources, which are summarized in **Supplemental Table S9**. For the LD tag and SNP-TSS distance datasets, we used those provided in the GARFIELD package.

Genomic coordinates of the annotations based on the GRCh38 genome were converted to the GRCh37 positions using the liftOver program downloaded from the UCSC Genome Browser Store (https://genome-store.ucsc.edu/). Comparison of the distributions of QTLs in the promoters of target genes and those in non-target gene promoters across the groups of tn>eQTLs, tn+eQTLs, and e>tnQTLs was performed by two-tailed Fisher’s exact test. In the analysis of enrichment in predicted TF binding sites, we focused on monomeric TFs whose expression was detected in one or more of the 17 cellular conditions (total *n* of TFs = 526). Functional elements with a standard error of ln(odds ratio) greater than 10 in the output result file were considered low confidence and were excluded from the visualization.

### Characterization of tn>eQTL-depleted TF binding regions

Protein-protein interactions among the 12 TFs (CUX2, FOXA1, FOXO6, LMX1A, PHOX2A, POU4F1, SOX1, SOX2, SOX3, SOX9, SRY, and ZNF211) whose binding regions were specifically depleted for tn>eQTLs was delineated by using STRING^77^ (version 12.0) with the following parameters: Network type = “full STRING network”; meaning of network edges = “confidence”; active interaction sources = “Textminging, Experiments, Databases, Co-expression, Neighborhood, Gene Fusion, and Co-occurrence”; minimum required interaction score = “low confidence (0.150)”. We used Cytoscape (version 3.10.2) for the network visualization. ZNF211 was not shown in the network figure because there was no inferred interaction with the other 11 TFs. GO enrichment analysis of these 12 TFs was performed by the *enrichGO()* function of clusterProfiler^24^ with the list of all expressed TFs as the background.

### DAF and conservation of QTLs and non-QTL SNPs

Determination of the ancestral and derived alleles of tn>eQTL, tn+eQTL, e>tnQTL, and the other non-QTL SNPs and the annotation of DAFs in the European population were performed using the 1000 Genomes Project reference data. For SNPs without derived allele information, we used MAFs instead. The conservation scores across 100 vertebrate species (phyloP100way) in the hg19/GRCh37 genome were downloaded from the UCSC GoldenPath (http://hgdownload.cse.ucsc.edu/goldenpath/hg19/phyloP100way/hg19.100way.phyloP100way.bw). The phyloP scores at the SNP positions were retrieved using the bigWigAverageOverBed tool. DAFs or phyloP100way scores across the four groups of SNPs were compared by the Mann-Whitney U test after performing LD-based pruning at r^2^ > 0.8 using plink (version 1.90b) with the following option: --indep-pairwise 100 5 0.8.

### Overlap between QTL target and constrained genes

The information on gene constraint metrics was obtained from the gnomAD (version 4.1) download site (https://storage.googleapis.com/gcp-public-data--gnomad/release/4.1/constraint/gnomad.v4.1.constraint_metrics.tsv). We considered canonical Ensembl genes whose transcript with “lof.oe_ci.upper” (loss-of-function observed/expected upper bound fraction, LOEUF) < 0.6 as constrained genes, as recommended in the gnomAD help page. Overlap between constrained genes and tn>eGenes, tn+eGenes, or e>tnGenes was evaluated by hypergeometric tests with all expressed genes as background using the *phyper*() function in R. We performed two-tailed tests according to the direction of overlap by doubling the *P* values obtained.

### LDSC and MAGMA analysis

#### GWAS summary statistics

GWAS summary statistics for the following 21 human traits were downloaded from the corresponding repositories (see the URL section for details): attention deficit/hyperactivity disorder (ADHD)^78^, autism spectrum disorder (ASD)^79^, bipolar disorder (BD)^80^, insomnia^81^, major depressive disorder (MDD)^82^, obsessive-compulsive disorder (OCD)^83^, schizophrenia^51^, amyotrophic lateral sclerosis (ALS)^84^, Alzheimer’s disease (AD)^85^, Parkinson’s disease (PD)^86^, inflammatory bowel disease (IBD)^87,88^, multiple sclerosis (MS)^89^, rheumatoid arthritis (RA)^90^, systemic lupus erythematosus (SLE)^91^, type 2 diabetes mellitus^92^, body mass index (BMI)^93^, educational attainment^94^, height^95^, intelligence^96^, neuroticism^97^, subjective well-being^98^. Since the summary statistics for IBD, MS, RA, SLE, and type 2 diabetes mellitus were based on the GRCh38 coordinates, variant positions were mapped onto the GRCh37 genome using the UCSC tool liftOver as needed. Variants without an rsID were annotated with the IDs in dbSNP151 (https://hgdownload.cse.ucsc.edu/goldenpath/hg19/database/snp151.txt.gz) using bcftools *annotate* (version 1.9).

#### Stratified LDSC

We analyzed enrichment of trait heritability in tn>eQTLs, tn+eQTLs, and e>tnQTLs using stratified LD SCore regression^39,40^ (LDSC, version 1.0.1). The aforementioned summary statistics were converted to the .sumstats format by *munge_sumstats.py* keeping the HapMap3 SNPs. Annotation files for the tn>eQTLs, tn+eQTLs, and e>tnQTLs were prepared by *make_annot.py* specifying the genomic positions of these QTLs in the UCSF bed format. LD scores based on the 1000 Genomes Phase 3 European dataset were computed with the output .annot files using *ldsc.py –l2*. We then calculated partitioned heritability for enrichment analysis by using *ldsc.py –h2*. Trait heritability enrichment in tn>eQTLs, tn+eQTLs, and e>tnQTLs was evaluated by using one-tailed *P* values.

#### MAGMA

Enrichment of aggregated genetic association in genesets (i.e. tn>eGenes, tn+eGenes, and e>tnGenes) of interest was analyzed by using Multi-marker Analysis of GenoMic Annotation^41^ (MAGMA, version 1.10). The annotation step to map SNPs to genes was performed by *magma --annotate* with 35kb upstream and 10kb downstream windows, following previous studies^99,100^. We then performed a gene analysis to compute gene-level metrics using *magma --gene-model snp-wise=top.* We used the *snp-wise=top* model, which is documented to be most sensitive when only a small fraction of the SNPs in a gene show an association in the MAGMA manual. Using the obtained gene-level results, we performed gene-set analyses for tn>eGenes, tn+eGenes, and the e>tnGenes in the integrated dataset of 17 conditions using *magma --gene-results*.

### Mendelian randomization

We performed two-sample Mendelian randomization (MR) using the R package TwoSampleMR^42^ (version 0.5.7). The tnQTL or eQTL SNPs were used as instrumental variables to investigate genetically determined transcriptional noise or gene expression dysregulation whose causal effects on one or more of the 21 human traits subjected to the above LDSC and MAGMA analyses were supported. We separately analyzed the data from the 17 cellular conditions.

For the exposure GWAS (i.e., tnQTL or eQTL mapping), the following information was included in the input: number of human iPSC donors, lfsr, posterior mean of effect estimate, posterior standard error, chromosomal coordinate, rsID, effect allele, non-effect allele, effect allele frequency, and the target gene. For the outcome GWAS (i.e., GWAS for the 21 traits) the following was used for the input: numbers of cases and controls for binary traits and the total number for non-binary traits, *P* value, slope, standard error, chromosomal coordinate, rsID, effect allele, non-effect allele, and effect allele frequency. Variants with rsIDs and *P* values < 1 × 10^-5^ were used for the analysis. After loading inputs by *read_exposure_data()* and *read_outcome_data()*, we harmonized the effect and non-effect alleles between the exposure and outcome data and removed ambiguous palindromic variants with intermediate allele frequencies using *harmonise_data()*. The variants in LD within a 10 Mbp window were then clumped by *ld_clump()* of the R package ieugwasr (version 0.1.5) with default parameters using the European LD reference panel of the 1000 Genomes Project^101^. MR estimates were obtained for each of the clumped variants by *mr_singlesnp(single_method = “mr_wald_ratio”)*. Directional horizontal pleiotropy was examined by *mr_pleiotropy_test()*. Traits with the estimates with a *P* value for the Egger intercept < 0.05 were not included in the subsequent analysis. Through this process, we excluded OCD. The clumped results were used to count the numbers of putative causal tnQTLs/tnGenes and eQTLs/eGenes. While we included the clumping step in the above main analysis, we also performed an analysis without clumping to comprehensively explore tn>eQTL SNPs associated with transcriptional noise dysregulation causal for multiple traits. This was because in some cases such SNPs were removed during the clumping step, despite their significant association.

### Prediction of TF binding motif alteration by tn>eQTL SNPs

We analyzed the TF-binding motifs that are predicted to be disrupted or created by tn>eQTL SNPs associated with transcriptional noise dysregulation whose causal effects on multiple traits were indicated by MR using the R packages motifbreakR^102^ (version 2.8.0), with BSgenome (version 1.62.0), SNPlocs.Hsapiens.dbSNP144.GRCh37 (version 0.99.20), BSgenome.Hsapiens.UCSC.hg19 (version 1.4.3), and MotifDb (version 1.36.0). Briefly, after loading the chromosomal coordinates of these tn>eQTL SNPs by *snps.from.file()*, candidate TFs whose motifs are altered by the tn>eQTL SNPs were sought by *motifbreakR()* with the following parameters: *pwmList = query(MotifDb, ‘hsapiens’), threshold = 1e-3, filterp = T, method = “ic”, bkg = c(A = 0.25, C = 0.25, G = 0.25, T = 0.25)*. We retrieved prediction *P* values by *calculatePvalue()*, and extracted TFs that were expressed in the cellular condition(s) where the SNP of interest had a significant tn>eQTL effect. For visualization, we used MotifDb (version 1.36.0) and seqLogo (version 1.60.0) R packages.

### C4 haplotypes

*C4* haplotypes of the donors of the human iPSC lines were imputed from the local SNP genotypes following the imputec4 protocol^103^ (https://github.com/freeseek/imputec4), using bcftools (version 1.9), beagle (version 5.1), and the European reference panel of haplotypes of the major histocompatibility complex region (MHC_haplotypes_CEU_HapMap3_ref_panel.GRCh37.vcf.gz downloaded from https://personal.broadinstitute.org/giulio/panels/). This procedure produces a count table of the following *C4* haplotypes: AL-AL-1, AL-AL-2, AL-AL-3, AL-AL-BL, AL-AL-BS, AL-AS-BL, AL-AS-BS-BS, AL-BL-1, AL-BL-2, AL-BL-3, AL-BL-other, AL-BS-1, AL-BS-2, AL-BS-3, AL-BS-4, AL-BS-5, AL-BS-BS, AL-BS-other, AL, BL and BS. We did not use the information of the minor haplotypes whose association with schizophrenia and SLE risks was not determined (AL-AL-3, AL-AL-BL, AL-AL-BS, AL-AS-BL, AL-AS-BS-BS, AL-BS-BS, AL and BL). The *C4* haplotype counts were subsequently summarized into the haplogroups^103^ (AL-AL = AL-AL-2 + AL-AL-1; AL-BL = AL-BL-3 + AL-BL-2 + AL-BL-1 + AL-BL-other; AL-BS = AL-BS-5 + AL-BS-other + AL-BS-4 + AL-BS-3 + AL-BS-2 + AL-BS-1; BS = BS) and their relationship with the rs4947249 genotypes were assessed.

### DEG and DNG analyses in schizophrenia patient and model brains

#### Human snRNA-seq data processing

We downloaded the processed data (gene × cell count matrices for each cell type, snRNA-seq_and_spatial_transcriptomics.zip) from a human schizophrenia case-control postmortem DLPFC (Brodmann area 9) snRNA-seq study^45^ from https://zenodo.org/record/6921620/files/. This dataset was derived from nine schizophrenia and 14 control brains and enriched for neuronal population sorted by neuronal marker NeuN. We used the cell type annotations assigned in the original study, consisting of eight excitatory (L2-CUX2-LAMP5, L2-3-CUX2-FREM3, L3-CUX2-PRSS12, L4-RORB-SCHLAP1, L4-5-FEZF2-LRRK1, L5-FEZF2-ADRA1A, L5-6-FEZF2-TLE4 and L5-6-THEMIS) and six inhibitory (ID2-LAMP5, ID2-NCKAP5, ID2-PAX6, PVALB, SST and VIP) types. Although some glial cells were included in this dataset, these cells were not used for our analysis because their numbers were small and they were not considered the primary focus of that postmortem brain snRNA-seq study. For each dataset of the 14 cell types, genes that were expressed (UMI count > 0) in < 1% of all cells (including both case and control cells) were excluded. We obtained denoised counts of gene expression for each of the 28 datasets (14 cell types × cases and controls) by correcting for the mean-variability dependency in gene expression using BASiCS^50^ (version 2.6.0), as described above. We included the sequencing instrument (“Novaseq_Instrument”; “CBMR” or “KU_FACS_core”) as a covariate in the BASiCS correction.

#### Mouse scRNA-seq data processing

The scRNA-seq data (bam files based on the mm10 mouse genome) from the PFC and striatum of two *Setd1a*^+/-^ and two wildtype (12-to 14-week-old) adult male mice^46^ were downloaded from the NCBI Sequence Read Archive (SRA, https://www.ncbi.nlm.nih.gov/sra) with the following accession numbers: SRR15278769 (PFC, wildtype, biological replicate 1), SRR15278770 (PFC, wildtype, biological replicate 2), SRR15278771 (PFC, *Setd1a*^+/-^, biological replicate 1), SRR15278772 (PFC, *Setd1a*^+/-^, biological replicate 2), SRR15278773 (striatum, wildtype, biological replicate 1), SRR15278774 (striatum, wildtype, biological replicate 2), SRR15278775 (striatum, *Setd1a*^+/-^, biological replicate 1), and SRR15278776 (striatum, *Setd1a*^+/-^, biological replicate 2). The cell-type information annotated in the Chen et al. study and the cell barcode IDs were retrieved from Gene Expression Omnibus (GEO) with the accession number GSE181021 (GSE181021_NAc_scRNAseq_meta.csv.gz, GSE181021_PFC_Excitatory_scRNAseq_meta.csv.gz and GSE181021_PFC_scRNAseq_meta.csv.gz). The following 17 cell types (ten in PFC and seven in striatum) were included in this dataset: excitatory neurons, inhibitory neurons, endothelial cells, astrocytes, microglial cells, oligodendrocytes, and oligodendrocyte precursor cells (OPCs) in the PFC, and *Drd1*-expressing medium spiny neurons (D1 MSN), *Drd2*-expressing MSN (D2 MSN), inhibitory neurons not expressing *Drd1* and *Drd2* (non-DRD inhibitory), endothelial cells, astrocytes, microglial cells, oligodendrocytes, OPCs, newly formed oligodendrocytes (NFOs), and neural stem cells/neuroblasts (NBs) in the striatum. Filtered gene × cell barcode matrices for each cell type per genotype (34 datasets in total) were produced from the above-described bam files following the analysis guide on the 10x Genomics webpage (https://www.10xgenomics.com/resources/analysis-guides/tutorial-navigating-10x-barcoded-bam-files). For each of these 34 datasets, we excluded genes expressed in < 1% of the cells and then corrected for mean-variability dependency using BASiCS^50^, as was performed for the human snRNA-seq data. We used mouse gene symbols and Ensembl gene IDs version 102 downloaded from Ensembl bioMart for annotation. Mouse orthologs of human genes were identified based on the human-mouse connection table provided by the Mouse Genome Informatics database requiring the exact matches of the gene symbols (https://www.informatics.jax.org/downloads/reports/HMD_HumanPhenotype.rpt).

#### DNG and DEG detection

Differentially noisy genes (DNGs) and differentially expressed genes (DEGs) between schizophrenia cases and controls or *Setd1a*^+/-^ and wildtype mice were identified by using the module for differential variability or mean test between two cell populations in the BASiCS^50^ (version 2.6.0) package, according to the developer’s instruction (https://bioconductor.org/packages/release/bioc/vignettes/BASiCS/inst/doc/BASiCS.html#analysis-for-two-groups-of-cells). The human and mouse datasets of variability (residual over-dispersion, a measure of how greater the variance is than expected, which is not affected by the mean) or mean of each gene in a cell population were analyzed by running the *BASiCS_TestDE()* function with the default parameters. This analysis with the default settings (EpsilonM = EpsilonD = log_2_(1.5)) identifies genes with differential residual over-dispersion (i.e. DNGs) or differential mean (i.e. DEGs) between the two groups with an absolute increase of 50% or more at the expected FDR of 0.05. We analyzed each cell type separately.

#### DNG and DEG characterization

Correlations across cell types were assessed by calculating pairwise Pearson’s rho for each gene’s case-control fold change in residual overdispersion or mean. In this analysis, outlier genes (< the first quartile - 1.5 × the interquartile range or > the third quartile + 1.5 × the interquartile range) were excluded. Stratified LDSC and MAGMA analyses were performed as described above, using DNGs or DEGs in each cell type as inputs instead of the QTL SNPs or their target genes. We note that we used the *--gene-model snp-wise=mean* option in the MAGMA analysis of DNGs and DEGs, considering that the numbers of input genes were in general larger than in the analysis of tn>eQTL target genes. We separately analyzed up- and down-regulated genes. General GO enrichment analysis of DNG or DEG sets of interest was performed by using the e*nrichGO()* function of clusterProfiler^24^ (version 4.2.2) as described above. An enrichment analysis focusing on synaptic genes was performed by using the SynGO^59^ web browser (version 1.2, https://www.syngoportal.org/) “Your gene list” module with all genes expressed in L2-CUX2-LAMP5 or D2 MSN as the background.

### Simulation of statistical power for QTL detection

We used powerEQTL^61^ (https://bwhbioinfo.shinyapps.io/powerEQTL/) to simulate the statistical power for QTL detection using scRNA-seq data. We evaluated the following three conditions: 1) Total numbers of subjects = 155 and Number of cells from each subject (m) = 357 (795,661 cells/17 conditions/131 donors on average), which correspond to the current study; 2) Total numbers of subjects = 300 and Number of cells from each subject (m) = 357, which simulate a larger study with the same per individual cell number; 3) Total numbers of subjects = 53 and Number of cells from each subject (m) = 106 (5,597 cells/53 donors), which correspond to the sample and cell sizes in the Sarkar et al. study. Other parameters were set to the default values (Slope under alternative hypothesis (β1) = 0.13; Standard deviation of gene expression (σ_y_) = 0.13; Intra-class correlation (range [0, 1]; ρ) = 0.5; Family-wise type I error rate (FWER) = 0.05; Total number of all (SNP, gene) pairs (nTests) = 20000), which were also employed in the PsychENCODE2 study^104^.

## Resource availability

### Lead contact

Requests for further information and resources should be directed to and will be fulfilled by the lead contact, Atsushi Takata.

### Materials availability

This study did not generate new unique reagents.

### Data and code availability

GENCODE comprehensive annotation v38lift37^33^,

https://ftp.ebi.ac.uk/pub/databases/gencode/Gencode_human/release_38/GRCh37_mapping/gencode.v38lift37.annotation.gtf.gz.

Ensembl bioMart, https://asia.ensembl.org/info/data/biomart/index.html.

SHAPEIT^72^, https://mathgen.stats.ox.ac.uk/genetics_software/shapeit/shapeit.html.

IMPUTE2^73^, https://mathgen.stats.ox.ac.uk/impute/impute_v2.html.

Reference panel of the 1000 Genomes Project Phase 3 for SHAPEIT and IMPUTE2, https://mathgen.stats.ox.ac.uk/impute/1000GP_Phase3.html.

FANTOM5 annotations^28,30,31^, https://fantom.gsc.riken.jp/5/datafiles/latest/extra/CAGE_peaks_annotation/DPIcluster_hg19_20120116.permissive_set.TATA_CpG_annotated.osc.gz,

https://fantom.gsc.riken.jp/5/datafiles/latest/extra/Enhancers/human_permissive_enhancers_phase_1_and_2.bed.gz,

https://fantom.gsc.riken.jp/5/datafiles/latest/extra/CAGE_peaks/hg19.cage_peak_phase1and2combined_ann.txt.gz.

UCSC Table Browser, https://genome.ucsc.edu/cgi-bin/hgTables.

JASPAR 2022 CORE collection^105^, http://hgdownload.soe.ucsc.edu/gbdb/hg19/jaspar/JASPAR2022.bb.

Human ancestral allele information, ftp://ftp.ensembl.org/pub/release-75/fasta/ancestral_alleles/homo_sapiens_ancestor_GRCh37_e71.tar.bz2.

phyloP score^106^, http://hgdownload.cse.ucsc.edu/goldenpath/hg19/phyloP100way/hg19.100way.phyloP100way.bw.

Gene constraint, https://storage.googleapis.com/gcp-public-data--gnomad/release/4.1/constraint/gnomad.v4.1.constraint_metrics.tsv.baseline-LD model v2.2, https://zenodo.org/records/10515792/files/1000G_Phase3_baselineLD_v2.2_ldscores.tgz.

GWAS summary statistics: attention deficit/hyperactivity disorder^78^,

10.6084/m9.figshare.14671965/daner_adhd_meta_filtered_NA_iPSYCH23_PGC11_sig PCs_woSEX_2ell6sd_EUR_Neff_70.meta.gz; autism spectrum disorder^79^,

10.6084/m9.figshare.14671989/iPSYCH-PGC_ASD_Nov2017.gz; bipolar disorder^80^,

10.6084/m9.figshare.14102594/pgc-bip2021-all.vcf.tsv.gz; insomnia^81^,

https://ctg.cncr.nl/documents/p1651/Insomnia_sumstats_Jansenetal.txt.gz; major depressive disorder^82^,

https://datashare.ed.ac.uk/handle/10283/3203/PGC_UKB_depression_genome-wide.txt.gz; obsessive compulsive disorder^83^,

10.6084/m9.figshare.14672103/ocd_aug2017.gz; schizophrenia^51^,

10.6084/m9.figshare.19426775/PGC3_SCZ_wave3.primary.autosome.public.v3.vcf.tsv.g z; amyotrophic lateral sclerosis^84^,

http://ftp.ebi.ac.uk/pub/databases/gwas/summary_statistics/GCST90027001-GCST90028000/GCST90027164/GCST90027164_buildGRCh37.tsv.gz; Alzheimer’s disease^85^,

https://ctg.cncr.nl/documents/p1651/AD_sumstats_Jansenetal_2019sept.txt.gz; Parkinson’s disease^86^,

https://drive.google.com/file/d/1FZ9UL99LAqyWnyNBxxlx6qOUlfAnublN/view/nallsEtAl2019_excluding23andMe_allVariants.tab.zip; inflammatory bowel disease^87,88^,

http://ftp.ebi.ac.uk/pub/databases/gwas/summary_statistics/GCST003001-GCST004000/GCST003043/harmonised/26192919-GCST003043-EFO_0003767.h.tsv.gz; multiple sclerosis^89^,

http://ftp.ebi.ac.uk/pub/databases/gwas/summary_statistics/GCST005001-GCST006000/GCST005531/harmonised/24076602-GCST005531-EFO_0003885.h.tsv.gz; rheumatoid arthritis^90^,

http://ftp.ebi.ac.uk/pub/databases/gwas/summary_statistics/GCST002001-GCST003000/GCST002318/harmonised/24390342-GCST002318-EFO_0000685.h.tsv.gz; systemic lupus erythematosus^91^,

http://ftp.ebi.ac.uk/pub/databases/gwas/summary_statistics/GCST003001-GCST004000/GCST003156/harmonised/26502338-GCST003156-EFO_0002690.h.tsv.gz; type 2 diabetes mellitus^92^,

http://ftp.ebi.ac.uk/pub/databases/gwas/summary_statistics/GCST006001-GCST007000/GCST006867/harmonised/30054458-GCST006867-EFO_0001360.h.tsv.gz; body mass index^93^,

https://portals.broadinstitute.org/collaboration/giant/images/c/c8/Meta-analysis_Locke_et_al%2BUKBiobank_2018_UPDATED.txt.gz; educational attainment^94^,

https://www.thessgac.com/papers/9/GWAS_EA_excl23andMe.txt.gz; height^95^,

https://portals.broadinstitute.org/collaboration/giant/images/f/f7/GIANT_HEIGHT_YENGO_2022_GWAS_SUMMARY_STATS_EUR.gz; intelligence^96^,

https://ctg.cncr.nl/documents/p1651/SavageJansen_IntMeta_sumstats.zip; neuroticism^97^,

https://ctg.cncr.nl/documents/p1651/sumstats_neuroticism_ctg_format.txt.gz; subjective well-being^98^,

https://www.thessgac.com/papers/9/SWB_Full.txt.gz.

LD block boundaries of the European population, https://bitbucket.org/nygcresearch/ldetect-data/src/master/fourier_ls-all.bed.

The European reference panel of haplotypes of the major histocompatibility complex region,

https://personal.broadinstitute.org/giulio/panels/MHC_haplotypes_CEU_HapMap3_ref_panel.GRCh37.vcf.gz. powerEQTL^61^,

https://bwhbioinfo.shinyapps.io/powerEQTL/.

Original codes used in this study will be available in a public repository.

## Supplemental information

Data S1-S2, Figures S1–S13 and Table S1-S9

Data S1. List of significant tnQTLs

Data S2. List of significant eQTLs

Figure S1. Adjustment for mean-variability dependency using BASiCS

Figure S2. GSEA of genes with large and small transcriptional noise

Figure S3. Genotyping data processing and quality controls

Figure S4. Statistics of tnQTL- and eQTL-gene pairs called by FastQTL

Figure S5. Overlap between tnQTLs/tnGenes and eQTLs/eGenes

Figure S6. Maximum and median expression levels of each gene across 17 cellular conditions

Figure S7. GO enrichment analysis of genes targeted by rotenone-suppressed or promoted tn>eQTLs

Figure S8. Enrichment analysis of tn>eQTLs, tn+eQTLs, and e>tnQTLs in repetitive elements

Figure S9. Characterization of QTL SNPs and their target genes with respect to evolutionary constraints

Figure S10. Additional local plots of tn>eQTLs associated with transcriptional noise dysregulation causal for multiple traits

Figure S11. The rs4947249 tn>eQTL SNP genotype frequencies stratified by *C4* haplogroups

Figure S12. Synaptic GO enrichment analysis of DNGs and DEGs in L2-CUX2-LAMP5 and D2 MSN

Figure S13. Statistical power for QTL detection from scRNA-seq data in different scenarios

Table S1. Statistics of scRNA-seq data from human iPSC-derived midbrain cells in 17 conditions

Table S2. Statistics of significant tnQTLs, eQTLs, tn>eQTLs, tn+eQTLs, and e>tnQTLs

Table S3. Enrichment analysis of tn>eQTLs, tn+eQTLs, and e>tnQTLs in functional genomic elements

Table S4. Results of LDSC and MAGMA analyses of tn>eQTLs, tn+eQTLs, and e>tnQTLs and their target genes

Table S5. Results of Mendelian randomization analysis of transcriptional noise and expression abundance

Table S6. Lists of DNGs and DEGs

Table S7. Results of LDSC and MAGMA analyses of DNGs and DEGs

Table S8. Sources of genotyping data

Table S9. Sources of functional genomic annotations

